# Multi-Task Brain Network Reconfiguration is Inversely Associated with Human Intelligence

**DOI:** 10.1101/2021.07.31.454563

**Authors:** Jonas A. Thiele, Joshua Faskowitz, Olaf Sporns, Kirsten Hilger

## Abstract

Intelligence describes the general cognitive ability level of a person. It is one of the most fundamental concepts in psychological science and is crucial for effective adaption of behavior to varying environmental demands. Changing external task demands have been shown to induce reconfiguration of functional brain networks. However, whether neural reconfiguration between different tasks is associated with intelligence has not yet been investigated. We used fMRI data from 812 subjects to show that higher scores of general intelligence are related to less brain network reconfiguration between resting state and seven different task states as well as to network reconfiguration between tasks. This association holds for all functional brain networks except the motor system, and replicates in two independent samples (*N* = 138, *N* = 184). Our findings suggest that the intrinsic network architecture of individuals with higher intelligence scores is closer to the network architecture as required by various cognitive demands. Multi-task brain network reconfiguration may, therefore, represent a neural reflection of the behavioral positive manifold – the essence of the concept of general intelligence. Finally, our results support neural efficiency theories of cognitive ability and reveal insights into human intelligence as an emergent property from a distributed multi-task brain network.

## Introduction

Intelligence captures the general cognitive ability level of a person. It is critically involved in learning from experiences and a prerequisite for effective adaption to changing environmental demands (Sternberg 1997). People who score high on tests of general intelligence perform better in multiple different cognitive tasks – an observation that is called the positive manifold of general intelligence (Spearman 1904). Although scientists started to investigate the biological underpinning of intelligence many decades ago, and correlates have been identified in brain structure (Gregory et al. 2016), brain function (Neubauer and Fink 2009), and in intrinsic brain connectivity (Hilger et al. 2017a, 2017b, 2020; for general reviews on neural correlates of intelligence see Jung and Haier 2007; Basten et al. 2015), it remains an open question whether there exists an equivalent of the positive manifold of general intelligence within the human brain, i.e., a ‘neuro-*g*’ (Haier 2017).

Intrinsic brain networks can be assessed in the absence of task demands during the so-called resting state (Biswal et al. 1995; Van den Heuvel and Hulshoff Pol 2010). Their topology has been suggested as reflection of information transfer between different brain regions and various topological network attributes have been related to differences in cognitive ability (Dubois et al. 2018; Hilger et al. 2020). Recently, the focus has broadened to include functional brain network interactions measured during active cognition, i.e., during task states (Braun et al. 2015; Cohen and D’Esposito 2016). Introducing such external task demands leads to task general and task-specific updates in functional connectivity (Cole et al. 2014) and was proposed to amplify relations between phenotypical variations and their neural basis, suggesting task-based connectivity as promising marker of general intelligence (Greene et al. 2018, 2020).

Brain network reconfiguration, defined as changes in fMRI-derived functional brain connectivity in adaption to different cognitive states, has previously been studied by comparing resting-state functional connectivity (i.e., intrinsic connectivity) with functional connectivity during tasks (Schultz and Cole 2016). Task-evoked changes in functional connectivity seem to be crucial for shifting neural processing (Cole et al. 2021), and the pioneering study of Schulz and Cole (2016) revealed a significant (negative) association between a global estimate of brain network reconfiguration and general intelligence. However, as the exact nature of changes has been shown to depend on the kind of task (Braun et al. 2015; Cohen and D’Esposito 2016; Soreq et al. 2021) as well as on the intensity level of the cognitive challenge (Shine et al. 2016; Hearne et al. 2017), considering brain network reconfiguration as a task-general phenomenon may only provide limited insights into underlying processes. More specific insights into general intelligence, i.e., into implicated cognitive processes and into a potential neural equivalent of the positive manifold, would therefore require the investigation of reconfiguration between different tasks. Such multi-task brain network reconfiguration has been demonstrated to capture meaningful variations between persons (Salehi et al. 2020) but has not yet been related to general intelligence. Furthermore, it has not yet been tested whether the association between brain network reconfiguration and general intelligence is driven by specific functional systems or whether it represents a whole-brain phenomenon. This would allow for additional insights about intelligence-relevant processes and how these processes are implemented on the neural level.

Here, we use fMRI data from a large sample of healthy adults (*N* = 812) assessed during different cognitive states, i.e., during resting state and during seven different task states, to test the hypothesis that higher levels of general intelligence relate to less brain network reconfiguration. Specifically, we expected this association to manifest in reaction to different cognitive demands and on various spatial scales. We used a straight-forward operationalization of brain network reconfiguration and implemented our analyses on a whole-brain level as well as on the level of seven and 17 canonical functional brain networks. The results confirm our hypotheses and suggest that functional brain networks of more intelligent people may require less adaption when switching between different cognitive states, thus, pointing towards the existence of an advantageous intrinsic brain network architecture. Further, we show that although the different cognitive states were induced by different demanding tasks, their relative contribution to the observed effect was nearly identical, a finding that supports the assumption of a task-general neural correlate, i.e., a neural positive manifold. Finally, the involvement of multiple brain networks suggest intelligence as an emergent property of a widely distributed multi-task brain network.

## Materials and Methods

### Participants

Main analyses were conducted on data from the HCP Young Adult Sample S1200 including 1200 Subjects of age 22-37 years (656 female, 1089 right-handed, mean age = 28.8 years). All study procedures were approved by the Washington University Institutional Review Board, and informed consent in accordance with the declaration of Helsinki was obtained from all participants (for details see Van Essen et al. 2013). Subjects with a Mini-Mental State Examination (MMSE) score ≤ 26 (serious cognitive impairment) or missing cognitive data needed for calculating a general intelligence factor were excluded. Cognitive measures of the remaining 1186 subjects were used as input for factor analysis to estimate a latent factor of general intelligence (see next section). After additional exclusion due to missing fMRI data and excessive head motion (see below), the final sample consists of 812 subjects (422 female, 739 right-handed, 22-37 years, mean age = 28.6 years).

### General intelligence *g*

To estimate a latent factor of general intelligence (*g*-factor), bi-factor analysis based on the Schmid-Leiman transformation (Schmid and Leiman 1957) was conducted in accordance to Dubois et al. (2018) for 12 cognitive measures (Table S1) of 1186 subjects.

### Data acquisition and preprocessing

We used fMRI data acquired during resting-state (four runs) and data acquired during seven tasks (two runs each) capturing information from eight different external demands, that are referred to as cognitive states in the subsequent text. Resting-state runs comprise 14:33 min data (1,200 time points), while task runs vary between 2:16 min (176 time points) and 5:01 min (405 time points) lengths. See Van Essen et al. (2013) for an overview of general data acquisition, Smith et al. (2013) for details of the resting-state acquisition, and Barch et al. (2013) for additional information about tasks. Briefly, all fMRI data was acquired with a gradient-echo EPI sequence (TR = 720 ms, TE = 33.1 ms, flip angle = 52°, 2 mm isotropic voxel resolution, multiband factor = 8) on a 3T Siemens Skyra with a 32-channel head coil. We used the minimally preprocessed HCP fMRI data (Glasser et al. 2013) and implemented further preprocessing comprising a nuisance regression strategy with 24 head motion parameters, eight mean signals from white matter and cerebrospinal fluid, and four global signals (Parkes et al. 2018). For task data, basis-set task regressors (Cole et al. 2019) were used simultaneously with the nuisance regressors to remove mean task-evoked neural activation. Finally, timeseries of neural activation were extracted from 200 nodes covering the entire cortex (Schaefer et al. 2018). In-scanner head motion was measured by framewise displacement (*FD*, Jenkinson et al. 2002). As recommended in Parkes et al. (2018), subjects were only included if mean *FD* < .2 mm, proportion of spikes (*FD* > .25 mm) < 20%, and no spikes above 5 mm were observed.

### Functional connectivity

Subject-specific weighted functional connectivity matrices (FC) were computed using Fisher z-transformed Pearson correlations between time series of neural activation from 200 cortical regions. For each of the eight states (rest, seven tasks), FC was first computed for RL and LR phase directions separately and averaged afterwards. Functional connections were then filtered based on their correlation with intelligence (*p* < .1, Finn et al. 2015; Shen et al. 2017). Connections inconsistently correlated with intelligence across states (positive in one, negative in another or vice versa) were excluded. To prevent circularity, this connection filtering step was cross-validated: First, the sample was divided into ten subsamples (by ensuring absence of family relations and equal distributions of intelligence scores via stratified folds). Second, intelligence-relevant connections (significantly correlated with intelligence, *p* < .1) were selected in nine subsamples only. And third, this selection of connections was then applied to the withheld subsample, thus in no case the correlation between functional connectivity and intelligence was calculated and applied in one and the same sample. Note, that such a filtering step has been applied in previous work to identify relations between functional connectivity and different phenotypical variations (e.g., Finn et al. 2015; Shen et al. 2017; Greene et al. 2018; Gao et al. 2019; Avery et al. 2020). Reconfiguration measures were calculated on a whole-brain level, as well as within and between pairs of networks based on the Yeo 7/17 canonical systems (Yeo et al. 2011). Note that for the analysis on the level of 17 functional networks, the *p*-threshold was increased to *p* < .2 to ensure a sufficient number of remaining connections (see Fig. 1 for a schematic illustration of the general workflow, Fig. S1 for the filtering procedure as well as Fig. S2 and Fig. S3 for an overview of the filtered and remaining functional brain connections).

**Fig. 1.**
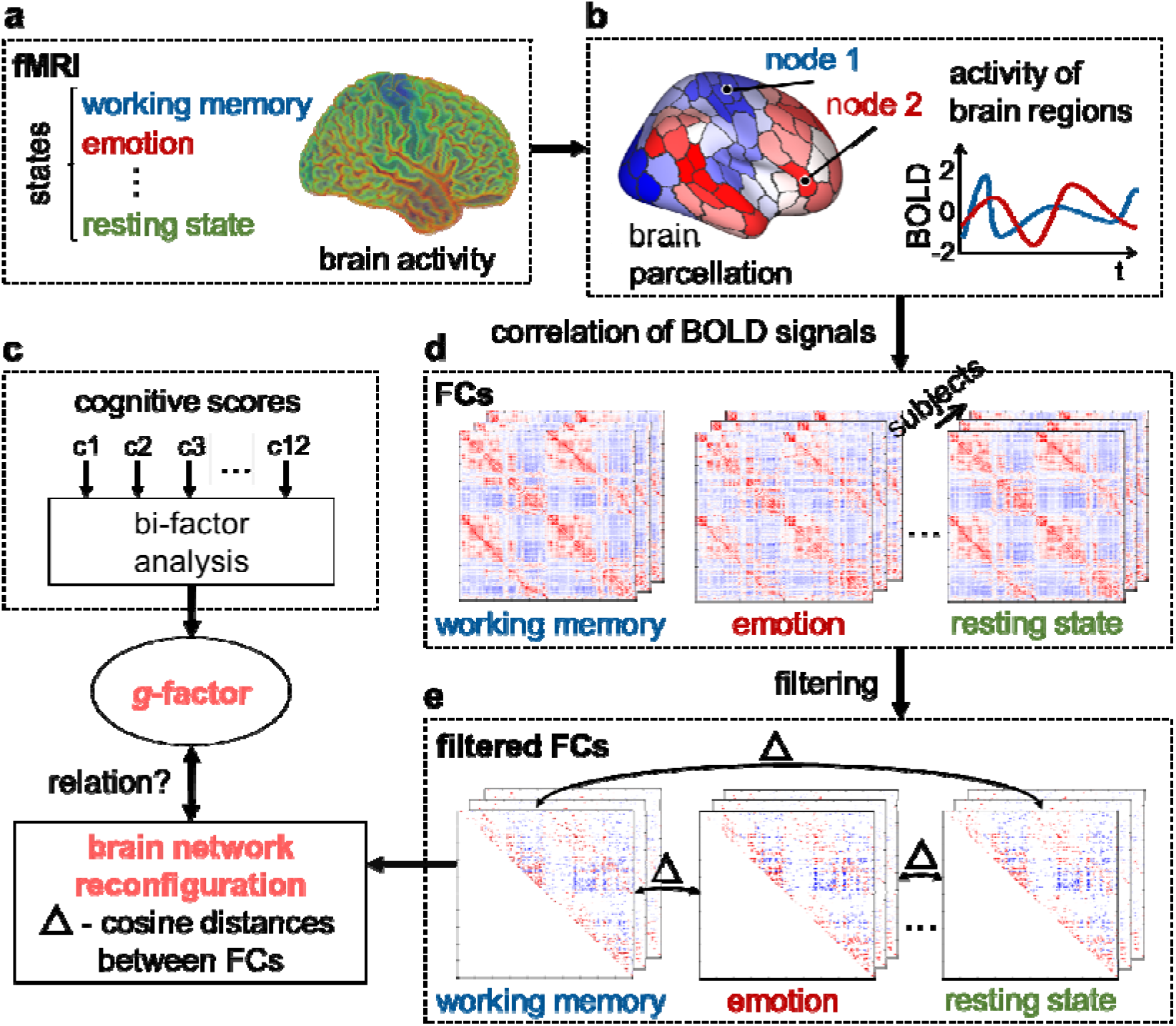
Schematic illustration of global analysis workflow. (**a**) Brain activity was assessed with fMRI during eight different cognitive states (resting state and seven tasks). (**b**) For each state, functional brain connectivity matrices (FCs, **d**) were computed by correlating the time series of 200 nodes with each other. For noise reduction, FCs were filtered based on their correlation with intelligence (see Fig. S1 and Methods for details). Brain network reconfiguration was calculated for all state combinations as cosine distances (Δ) between the filtered FCs (**e**). To assess the relationship between brain network reconfiguration and intelligence, reconfiguration values were correlated (Spearman correlations, controlled for age, sex, handedness and in-scanner head motion) with a latent *g*-factor derived from 12 cognitive scores using a bi-factor analysis model (**c**). BOLD, Blood oxygen level dependent (signal); t, time; c, cognitive score.

### Brain network reconfiguration

Reconfiguration of functional connectivity was operationalized as cosine distance between the filtered FCs of two states. The cosine distance *d_cos_* is the complement of the cosine similarity *s_cos_*:

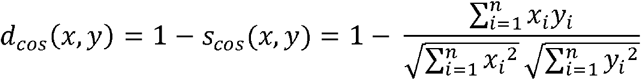

Where *s_cos_* is the cosine of the angle between two connection weights vectors *x* and *y* with a total number of connections *n*, which is expressed as the normalized inner product of the vectors. Note, that the cosine distance captures changes in orientation between two vectors and thus, indexes changes in the architecture (structure) of functional connectivity rather than changes in the strengths of connections (as captured with, e.g., Manhattan distance, Euclidean distance).

### Association between reconfiguration and intelligence

Relations between reconfigurations and intelligence were assessed with Spearman rankorder partial correlations by controlling for age, sex, handedness, and in-scanner head motion (mean *FD* over all scans and mean of percentage of spikes > .25 mm over all scans). For multiple comparisons, *p*-values were FDR corrected (*α* = .05). For gaining comprehensive insights not only into the general relation between brain network reconfiguration and intelligence, but also into the relevance of different states and the contribution of different brain networks, multiple reconfigurations values were computed for each participant: A) An average score of whole-brain reconfiguration between resting state and all task states (rest-task reconfiguration). B) An average score of whole-brain reconfiguration between all pairs of task states (task-task reconfiguration). C) 28 scores capturing whole-brain reconfiguration for each pair of rest-task and task-task state combinations (state combination-specific reconfiguration). D) Eight scores capturing wholebrain reconfiguration associated with one specific cognitive state (average over all 28 state combinations a specific state was involved in, i.e., state-specific reconfiguration). Note that for task states, only combinations with different tasks (no rest) were included. E) Brain-network specific reconfigurations scores for seven (and 17) functional brain networks (reconfigurations of all within- and between network combinations for each state combination). For interpretable insights, these network-specific reconfiguration scores (case E) were averaged a) over all state combinations (resulting in 28 (153) reconfiguration scores specific to a certain brain network combination), b) over all state combinations, and over all network-combinations the respective network was involved in (resulting in seven (17) state-independent network-specific reconfiguration scores), and c) over all state combinations a respective state was involved in (for task states, only combinations with different tasks were included), and over all seven (17) network combinations a respective network was involved in, in total summing up to 7 (17) network-specific reconfiguration scores for each state.

### External replication

For testing the robustness of our findings against varying measures of intelligence, varying cognitive demands induced by different tasks, and sample dependence, all analyses were repeated in two independent data sets (PIOP1, PIOP2) from The Amsterdam Open MRI Collection (AOMIC, Snoek et al. 2021). All study procedures were approved by the faculty’s ethical committee before data collection started (PIOP1 EC number: 2015-EXT-4366, PIOP2 EC number: 2017-EXT-7568) and informed consent in accordance with the declaration of Helsinki was obtained from all participants (for more details see Snoek et al. 2021). PIOP1 includes fMRI data of 216 subjects collected from six cognitive states (resting state and five tasks: emotion matching, gender-stroop, working memory, face perception, and anticipation), while PIOP2 contains fMRI data of 226 subjects from four states (resting state and three tasks: emotion matching, working memory, stop signal). Details on imaging parameters are described in Snoek et al. (2021). In brief, all fMRI data was acquired with a gradient-echo EPI on a Philips 3T scanner with a 32-channel coil (3 mm isotropic voxel resolution). Multiband scans were acquired for the face perception and resting-state paradigms of the PIOP2 sample (TR = 750 ms, TE = 28 ms, flip angle = 60°, multiband factor = 3). Sequential scans were acquired for resting state of the PIOP 2 sample, and the working memory, emotion matching, gender-stroop, anticipation, and stop signal paradigms of the PIOP1 and PIOP2 samples (TR = 2000 ms, TE = 28 ms, flip angle = 76.1°).

The Raven’s Advanced Progressive Matrices Test (36 item version - set II, Raven et al. 1998) was used in both samples for measuring intelligence. After excluding subjects with missing descriptive and behavioral data and after applying motion exclusion criteria (see above), 138 subjects (PIOP1) and 184 subjects (PIOP2) remained for analyses. The fMRI data was downloaded in the minimal preprocessed form, using an alternative preprocessing pipeline (fMRIprep v1.4.1, Esteban et al. 2019). Further preprocessing to extract nuisance regressed time series followed the same steps as specified above. As the PIOP samples are relatively small compared to the main sample and brain-behavior relationships are less reliable in small samples (Assem et al. 2020; Marek et al. 2020), no *p*-threshold was used for the selection of functional connections here. Instead, and to increase the robustness of this analyses, a filter mask was computed from the larger main sample (containing connections correlating only either positively or negatively with intelligence *p* < .01 in at least one of the filtered FCs of intersecting state combinations) and only connections located in these main sample filter mask and correlating with intelligence in the same direction in the replication samples were used in analyses.

### Data and Code availability

All analysis code used in the current study was made available by the authors on GitHub: Preprocessing: https://github.com/faskowit/app-fmri-2-mat; Main analyses: https://github.com/jonasAthiele/BrainReconfiguration_Intelligence, https://doi.org/10.5281/zenodo.5031683. All data used in the current study can be accessed online under: https://www.humanconnectome.org/study/hcp-young-adult (HCP), https://doi.org/10.18112/openneuro.ds002785.v2.0.0 (AOMIC-PIOP1), and https://doi.org/10.18112/openneuro.ds002790.v2.0.0 (AOMIC-PIOP2).

## Results

### Intelligence

General intelligence was operationalized as latent *g*-factor from 12 cognitive measures (Table S1) computed with bi-factor analysis (Dubois et al. 2018) using data from 1186 subjects of the Human Connectome Project (Van Essen et al. 2013). As per model fit criteria of Hu & Bentler (1999), the 4-bi-factor model fits the data well (Comparative Fit Index: CFI = .979, Root Mean Square Error of Approximation: RMSEA = .0395, Standardized Root Mean Square Residual: SRMR = .0213). The statistical model and the *g*-factor distribution in contrast to the PMAT-score distribution (brief assessment of intelligence provided by the HCP) is shown in Fig. S4.

### Less brain network reconfiguration is associated with higher intelligence

Brain network reconfiguration was operationalized as cosine distance between filtered functional connectivity matrices (FCs) of two out of eight different cognitive states (see Methods, Fig. 1 for a schematic illustration of the analyses workflow, Fig. S1 for details about the FC filtering procedure). Averaged across all rest-task and task-task state combinations, less brain network reconfiguration was associated with higher intelligence scores (rest-task: *rho* = -.23, *p* < .001; task-task: *rho* = -.23, *p* < .001; Fig. 2a). This effect also holds when using stricter thresholds for the cross-validated filtering approach, e.g., *p* < .01 (Table S2) or when using alternative mathematical operationalizations of reconfiguration (Pearson correlation between Fisher z-transformed FCs: rest-task: *rho* = .23, *p* < .001, task-task: *rho* = .23, *p* < .001; Manhattan distance between bi-partitioned FCs: *rho* = -.19, *p* < .001, task-task: *rho* = -.24, *p* < .001).

**Fig. 2.**
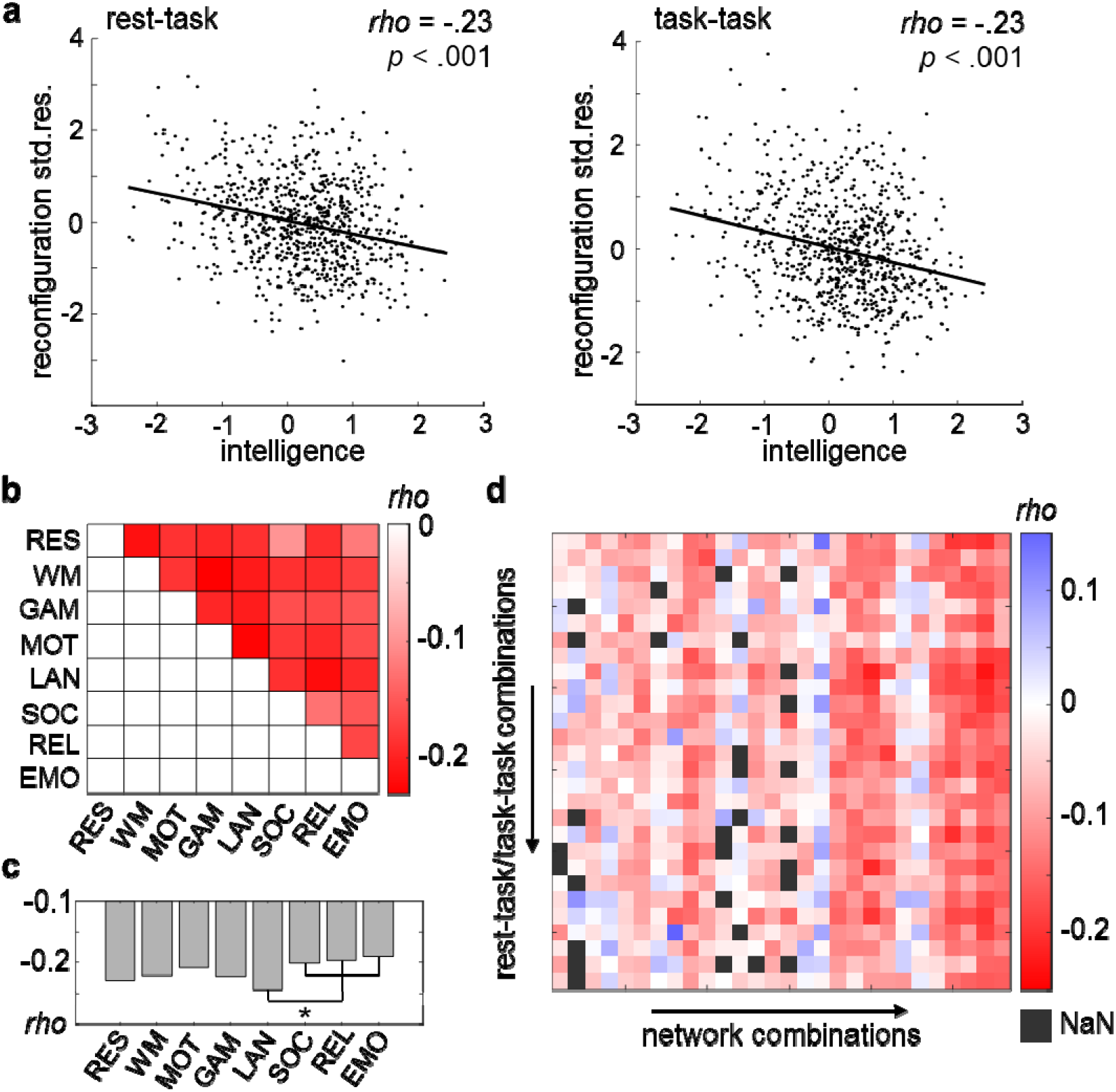
Less brain network reconfiguration is associated with higher intelligence. (**a**) Scatterplots illustrating the association (partial Spearman correlation, *rho*) between a latent *g*-factor of intelligence (derived from 12 cognitive tasks, *x*-axis) and the standardized residuals resulting from linear regression of age, sex, handedness, and in-scanner head motion on the variable of interest, that is brain network reconfiguration. Brain network reconfiguration was operationalized as cosine distance between functional connectivity matrices averaged over all possible rest-task combinations (*y*-axis, left panel), and all possible task-task combinations (*y*-axis, right panel) respectively. Note that only in this subfigure data of one subject was excluded due to visualization purposes. (**b**) Association between intelligence and brain network reconfiguration for all possible state combinations (FDR-corrected *p*-values, *α* = .05, all correlations are significant). The strengths of correlations are depicted in different colors (see color bar). (**c**) Associations between intelligence and a total measure of state-specific reconfiguration. i.e., reconfiguration values were averaged over all state combinations the respective state was involved in (FDR-corrected *p*-values, *α* = .05, all correlations are significant). Note that for task states, only combinations with different tasks (no rest) were included. Significant differences in correlation values (*p* < .05, marked with an asterisk) were only observed for the comparisons between the language task (LAN) and the social cognition, the relational processing, and the emotion processing task (SOC, REL, EMO) respectively. (**d**) Associations between intelligence and brain network- and state combination-specific reconfiguration values. Brain networks were derived from the Yeo atlas (Yeo et al., 2011, seven network partition used here) and network combinations refer to all within and between network connectivity combinations (columns). The strengths of correlations are depicted in different colors (see color bar). Note that NaN (not a number) values exist if in a specific network-state combination no single brain connection passes the filtering procedure (see Fig. S1 and Methods). For details about the assignment of the correlation values to the specific state and network combinations, see Fig. S5. Std. res., standardized residuals; FDR, false discovery rate; RES, resting state; WM, working memory task; GAM, gambling task; MOT, motor task; LAN, language processing task; SOC, social cognition task; REL, relational processing task; EMO, emotion processing task.

### Higher intelligence is related to less reconfiguration across different cognitive demands

Significant associations between higher intelligence and less brain network reconfiguration were observed for all rest-task and task-task state combinations (Fig. 2b). The correlations between reconfiguration and intelligence ranged from *rho* = -.10 (*p* = .006) for reconfiguration between resting state and social recognition task to *rho* = -.23 (*p* < .001) for reconfiguration between working memory and motor task. Again, similar associations were observed when using alternative reconfiguration metrics (Fig. S5). For evaluating the total influence of each individual state on the observed effect, reconfiguration values were averaged across all resttask combinations (for resting state) and separately over all task-task combinations a respective task was involved in (for each task state). The total influence of the language task was significantly stronger (*p* < .05) than the influence of the social recognition task, the relational processing, and the emotion processing task, while all other states did not differ significantly in their total influence on the observed effect (Fig. 2c and Fig. S6).

### The relation between reconfiguration and intelligence depends on different functional brain systems rather than on specific cognitive demands

By parcellating the brain into seven functional networks (Yeo et al. 2011), and by considering all possible network and state combinations, we observed that the variance of the effect between different state combinations was significantly smaller than the variance of the effect between different network combinations (Wilcoxon rank sum test, W = 441, *p* < .001, Fig. 2d, Fig. S7). This suggests prior importance of the differentiation between different brain systems rather than between different external demands.

### Higher intelligence is related to less reconfiguration across different spatial scales

Next, we analyzed the relative contribution of seven and 17 functional brain networks to the observed effect. Overall, higher intelligence scores were associated with less reconfiguration of within and between network connectivity in multiple brain networks. Dorsal and ventral attention systems, the control network, the default mode network, and limbic areas showed consistent significant negative associations, while in the visual and somatomotor networks the effect was weaker and the pattern more heterogeneous (Fig. 3a). To derive a more global measure of total network-specific reconfiguration, we then aggregated reconfiguration scores across all network-combinations a respective network was involved in. Higher intelligence was significantly associated with less connectivity reconfiguration in respect to all networks, except the somatomotor system (Fig. 3b). Similar relations were observed within and between 17 functional brain networks (Fig. 3c,d).

**Fig. 3.**
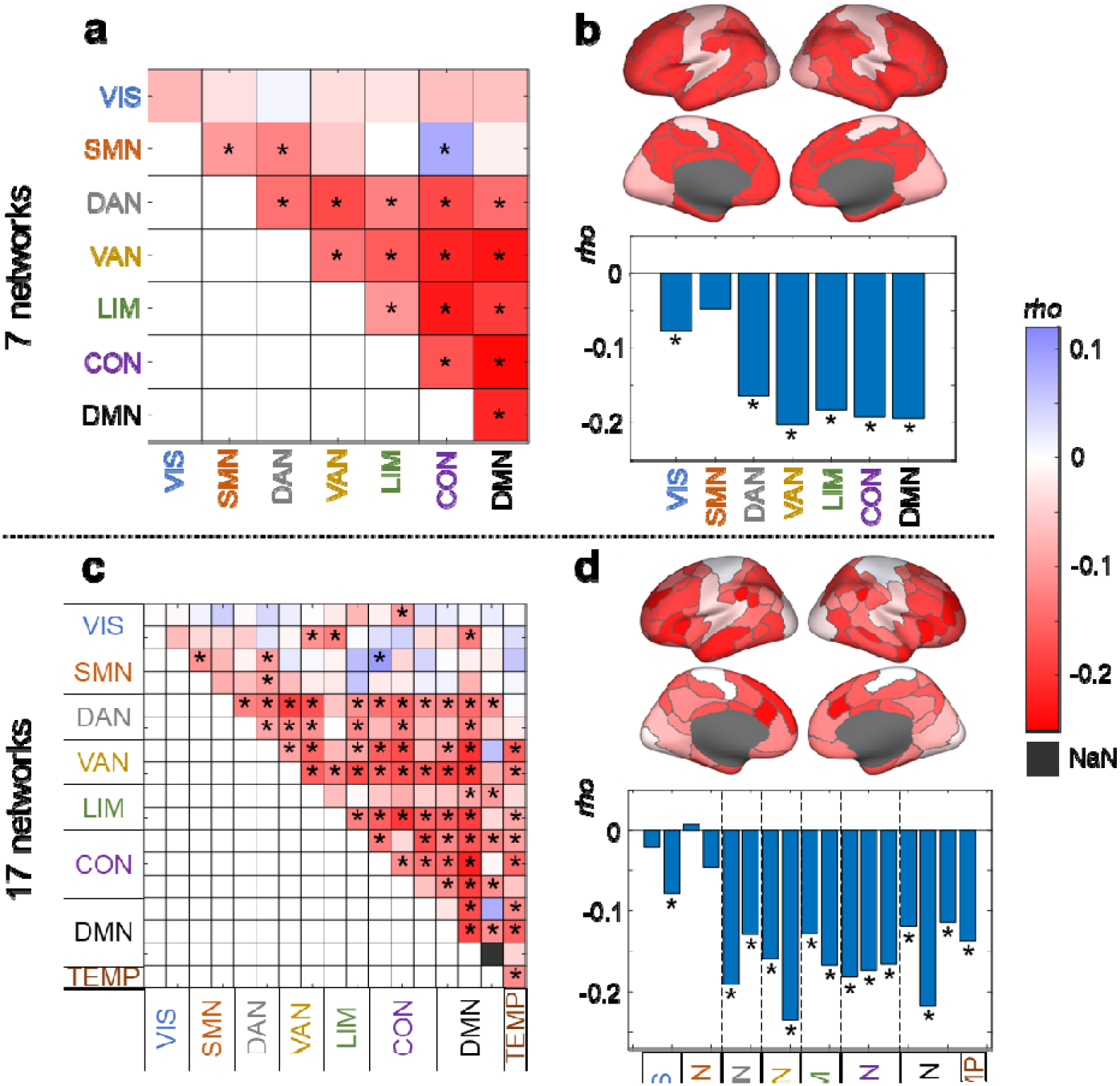
Brain network-specific associations between general intelligence and brain network reconfiguration. Partial spearman correlation (*rho*) between intelligence (*g*-factor derived from 12 cognitive tasks) and brain network-specific reconfiguration values (averaged cosine distance between functional connectivity matrices of eight different cognitive states) for seven and 17 separate functional brain networks (Yeo et al., 2011). Network-specific correlations were projected onto the surface of the brain. The strengths of correlations are depicted in different colors (see color bar). All correlations were controlled for influences of age, sex, handedness, and in-scanner head motion. All significant correlations (FDR-corrected *p*-values, *α* = .05) are marked with asterisks. Note that NaN (not a number) values exist if in a specific network combination no single brain connection passes the filtering procedure (see Fig. S1 and Methods). (**a**) Associations between intelligence and brain network combination-specific reconfiguration scores for seven functional brain networks. (**b**) Associations between intelligence and reconfiguration scores for seven functional brain networks (averaged across all within- and between network combinations a respective network is involved in). (**c**) Associations between intelligence and brain network combination-specific reconfiguration scores for 17 functional brain networks. (**d**) Associations between intelligence and reconfiguration scores for 17 functional brain networks (averaged across all within- and between network combinations a respective network is involved in). FDR, false discovery rate; VIS, visual network; SMN, somatomotor network; DAN, dorsal attention network; VAN, salience/ventral attention network; LIM, limbic network; CON, control network; DMN, default mode network; TEMP, temporal parietal network.

### Network-specific reconfigurations in response to varying external demands

Finally, we investigated network-specific contributions on the association between intelligence and brain network reconfiguration for each cognitive state. To this end, network-specific reconfiguration scores were aggregated across all rest-task combinations (for resting state) or task-task combinations a respective task was involved in (for each task state). As illustrated in Fig. 4, network-specific associations between reconfiguration and intelligence were relatively stable across all cognitive states. Again, similar relations were observed at the level of 17 functional brain networks (Fig. S8).

**Fig. 4.**
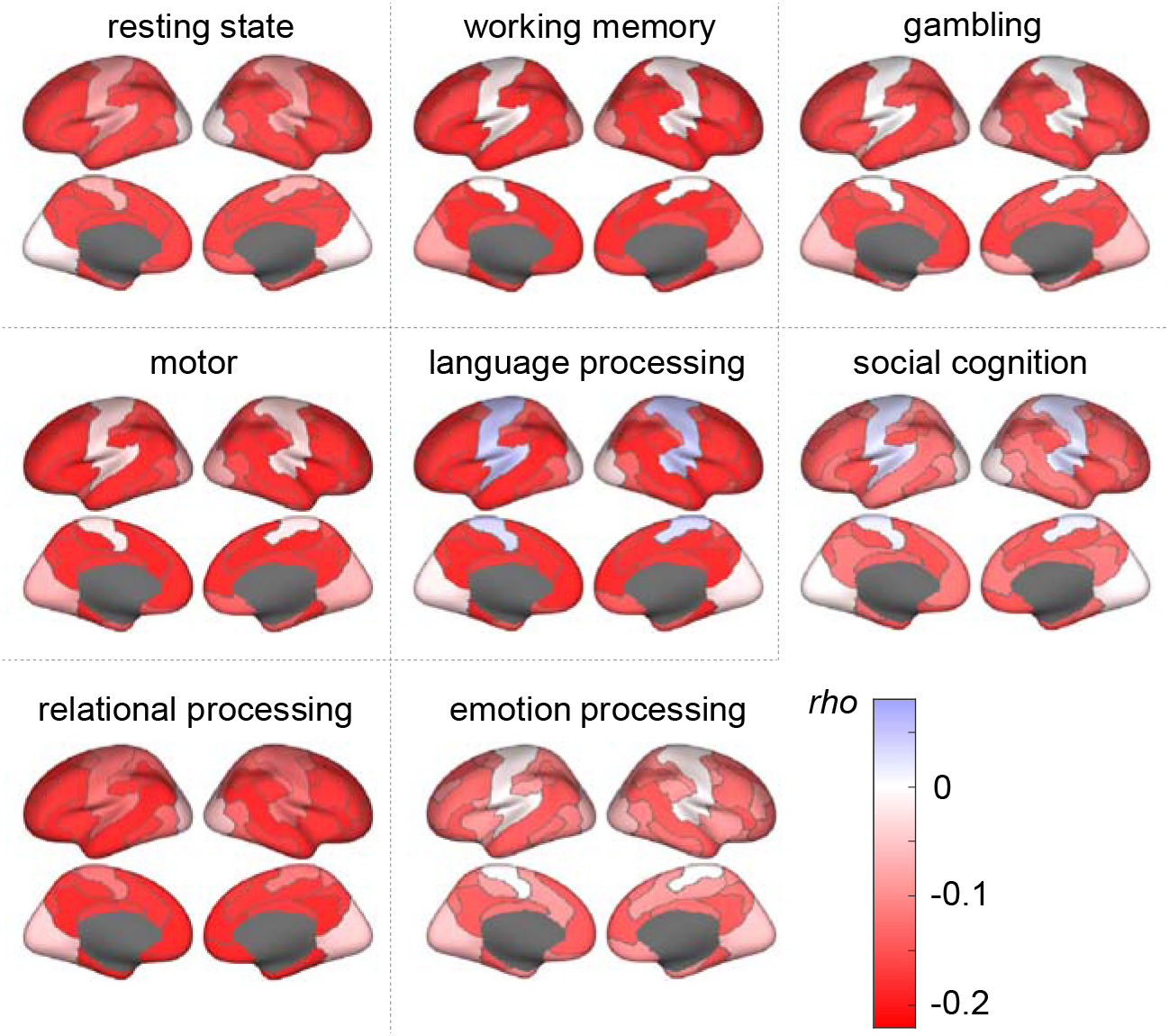
Associations between intelligence and brain network reconfiguration for each cognitive state. The strengths of partial Spearman correlations (*rho*) between intelligence (*g*-factor derived from 12 cognitive tasks) and brain network reconfiguration (cosine distance between functional connectivity matrices of different states) are illustrated in different colors (see color bar). All associations were controlled for influences of age, sex, handedness, and in-scanner head motion. Seven functional brain networks were derived from the Yeo atlas (Yeo et al., 2011). For calculating state-specific associations, cosine distances were averaged over all state combinations a respective state was involved in (for task states, only combinations with different tasks were included), and averaged over all network combinations (within- and between network connectivity) the respective network was involved in.

### External replication: Generalization to different measures of intelligence and different cognitive demands

To evaluate the robustness of our findings against different measures of intelligence and against varying cognitive demands induced by different tasks, all analyses were repeated in two independent samples (The Amsterdam Open MRI Collection AOMIC, Snoek et al. 2021, PIOP1: *N* = 138, PIOP2: *N* = 184, see Methods). In line with our main analyses, less brain network reconfiguration was associated with higher intelligence. This effect holds for both rest-task and task-task reconfiguration (PIOP1 rest-task: *rho* = -.32, *p* < .001, task-task: *rho* = -.26, *p* = .003; PIOP2 rest-task: *rho* = -.23, *p* = .002, task-task: *rho* = -.26, *p* < .001, Table S2) and became visible across most state combinations (Fig. S6). In PIOP1, nine out of 15 rest-task and task-task state combinations showed a significant negative association (range: -.12 ≤ rho ≤ -.35, .001 < *p* ≤ .18), while in PIOP2, five out of six rest-task and task-task state combinations showed the respective effect (range: -.13 ≤ *rho* ≤ -.30, .001 < *p* ≤ .09). Aggregating across all state combinations in which a respective state was involved in demonstrated that only in the PIOP2 sample the total influence of the stop signal task was significantly stronger (*p* < .05) than the total influence of the emotional matching task. All other states did not differ significantly in their total influence on the observed effect (Fig. S6). Finally, results from network-specific analysis were also similar to the results from the main sample (Fig. S9, S10). In sum, the results of the replication analyses support the robustness of our findings and suggest that the association between higher intelligence scores and less brain network reconfiguration generalizes to different cohorts, imaging acquisition parameters, operationalizations of intelligence, and to different cognitive demands.

### Robustness control analyses

Although the adopted procedure for filtering out noise-contaminated functional brain connections was thoroughly cross validated (see Methods) rendering potential circularity of analyses unlikely, to evaluate any remaining conceivable possibilities that results are biased by this step, all whole-brain analyses were repeated by a) considering all possible functional brain connections (i.e., no filter for selecting intelligence-related connections) and b) implementing a filter based on the pure overlap of intelligence-related edges and ignoring the sign of the association between connectivity and intelligence (different filter). The same association between higher intelligence and less brain network reconfiguration were observed in both cases (no filter: rest-task: *rho* = -.12, *p* < .001; task-task: *rho* = -.12, *p* < .001; different filter: rest-task: *rho* = -.21, *p* < .001; task-task: *rho* = -.21, *p* < .001). Without filtering, state-specific effects were overall smaller with not all state combinations reaching the significance threshold, while state-specific results based on the different filtering procedure were nearly identical (see Table S2, Fig. S11). Together, these analyses suggest that our filtering procedure successfully reduced noise and, most importantly, demonstrate that the observed association between higher intelligence and less brain network reconfiguration does not represent a spurious result of the filter.

To rule out that our results were influenced by region-specific or subject-specific differences in data quality (e.g., signal drop out), we repeated our main analysis while additionally controlling for brain subject- and brain network-specific differences in temporal signal to noise ratio (tSNR) of the minimal preprocessed BOLD signals (i.e., mean of BOLD signal divided by its standard deviation). With the additional control of whole-brain subject-specific tSNR values, the relation between intelligence and whole brain network reconfiguration was nearly identical to the relation resulting from the main analysis (with additional tSNR control: rest-task: *rho* = -.24, p < .001; task-task: *rho* = -.24, *p* < .001; without SNR control: rest-task: *rho* = -.23, *p* < .001; task-task: *rho* = -.23, *p* < .001). Also, the results of network-specific associations were highly similar to the main analysis when subject- and network-specific tSNR values were added as additional control variables (see Fig. S12). This demonstrates that our results were not substantially influenced by individual- or brain network-specific differences in tSNR.

Finally, to preclude the possibility that our findings were influenced by differences in scan duration (varying between 176 data points for the emotion task and 1200 data points for rest), all whole brain analyses were repeated while reducing all scans to the shortest scan duration (176 data points). Specifically, we selected 176 consecutive time points of each scan, starting from a randomly chosen time point. Reconfiguration of all rest-task and task-task state combinations as well as all aggregated state-specific reconfigurations were similarly related to intelligence as in the main analysis (for a detailed comparison see Fig. S13).

## Discussion

We showed that general intelligence is associated with less brain network reconfiguration expressed by higher similarity between functional brain connectivity linked to various cognitive states. In line with our initial hypothesis, this effect was not only observed for reconfiguration between rest and task but also for reconfiguration between different tasks each associated with a specific cognitive demand. Multiple control analyses and replication in two independent samples demonstrate the robustness of our findings and suggest generalizability of this effect to different measures of intelligence and to various cognitive demands. Finally, multiple functional brain systems were identified as driving this effect suggesting that intelligence is an emergent whole-brain phenomenon.

Our finding that less reconfiguration of functional brain connectivity is related to higher intelligence supports the assumption that more intelligent people may have an intrinsic brain network architecture that is better suited to fulfil multiple cognitive demands. In this regard, intrinsic brain connectivity, as assessed during rest, can be understood as baseline or inherent network architecture that undergoes task-specific adaptations to optimally support upcoming external demands (Cole et al. 2021). The observation that higher intelligence is not only associated with less reconfiguration between rest and task but also with less reconfiguration between different tasks suggests that the intelligence-associated advantage in network adaption primarily refers to task-general (in contrast to task-specific) adaptions, which have been shown to build the major proportion of functional connectivity changes when external demands are induced (Cole et al. 2014). On this basis, we speculate that an intrinsic network architecture which is closer to such a general task-supporting functional brain network structure may allow more intelligent people to switch faster and more efficiently (in terms of energy consumption) between rest and task as well as between different tasks associated with varying cognitive demands. Lower reaction times (Jensen 2006) as well as smaller latencies of event-related brain potentials in reactions to upcoming task stimuli (Schubert and Frischkorn 2020) have both been associated with higher intelligence. Future studies that relate these behavioral and electrophysiological measures to varying amounts of fMRI-derived network reconfiguration in one and the same sample may provide further insights into associations between those measures.

Further, our findings can also be interpreted against the background of fMRI task activation studies, in which less brain activation was associated with more efficient stimulus processing (Lustig and Buckner 2004; Liu and Pleskac 2011) and with higher intelligence (Neubauer and Fink 2009). Both were suggested as supporting the Neural Efficiency Hypothesis of intelligence that globally suggests that more intelligent people show less neural effort to adapt to a specific task (Haier et al. 1988; Dunst et al. 2014).

The observation that all tasks seem to contribute to the observed effect with almost equal strength may support the assumption that task-general adaptations have greater importance for intelligence-associated processing advantages than task-specific adaptations. The overlap of network architectures as required by different cognitive demands (i.e., intrinsic connectivity + task-general adaptions) can thus be interpreted as reflection of the positive manifold of general intelligence on the neural level (Spearman 1904; Kovacs and Conway 2016). In that, our study lend support to one of the oldest theories of human intelligence (*g*-factor, Spearman 1904) and provides at least a preliminary answer to the question about the existence of a ‘neuro-*g*’ (Haier 2017).

Moreover, we analyzed the impact of variations between different functional brain systems on the observed effect and showed that the relation between reconfiguration and intelligence depends more on variations between different brain networks than on variations between different task states. Specifically, reconfiguration in all brain systems except in the somatomotor network was significantly related to intelligence. The here proposed neural positive manifold, i.e., the overlap of network architectures as required by different cognitive demands, may thus include most but not all functional brain systems. However, although our replication supports the generalizability to an additional set of cognitive states, the selection of tasks was still limited in the current study and it therefore requires further investigation to test whether this effect is universal for a broader range of cognitive demands.

Overall, the involvement of multiple functional brain systems supports major neurocognitive models of intelligence like the Parieto-Frontal Integration Theory (P-FIT, Jung and Haier 2007), the Multiple Demand System Theory (MD, Duncan 2010) as well as meta-analytic findings (Basten et al. 2015; Santarnecchi et al. 2017, 2021) that reveal individual differences in intelligence to be not only associated with variations in structural or functional characteristics of a single brain region, but rather more to properties of a distributed network with major implications of neural systems associated with attentional control (Hilger et al. 2017a, 2020), executive functioning (Unsworth et al. 2009), and the default-mode of brain function (Basten et al. 2013). Although limbic brain systems have long been neglected by most of intelligence research, recent evidence supports these brain regions implication also in cognition (Catani et al. 2013) and our study can further contribute to this accumulating evidence.

Several limitations need to be mentioned. First, although we applied in-sample crossvalidation strategies to increase the generalizability of the functional connectivity filter, we cannot completely rule out any remaining influences of sample-specific characteristics on the connection selection procedure. This could impact functional connectivity results especially as we observed that filters became instable in smaller samples (replication samples). To address this issue, we conducted a conservative approach and applied the filter mask of the main sample (HCP) to both replication samples for increasing robustness. Future studies may take sample size into strong consideration and draw specific attention to construct robust and across-sample generalized functional connection selection strategies. Second, the sample of our study was restricted to subjects between 22-37 years of age; thus, future studies should address the question whether results generalize to a broader age range. Third, although our replication analysis shows generalizability of our finding to a different intelligence test and different cognitive demands, future studies may contribute to further broaden the picture to more diverse tasks as well as tasks that are more directly associated with the intelligence test assessment (Soreq et al. 2021) including multiple levels of difficulty (Dunst et al. 2014; Hearne et al. 2017; Sripada et al. 2020). Such investigations would be valuable for gaining more comprehensive insights into the role different cognitive processes may play within the relationship between brain network reconfiguration and general intelligence. Fourth, in contrast to task activation studies that attempt to identify the neural correlate of more specific circumscribed cognitive subprocesses during an cognitive task conducted in the scanner by calculating specific task contrasts (e.g., 2-back minus 0-back, Ragland et al. 2002; Blokland et al. 2008), we here compare complete tasks with each other and, thus, treated them as unified concepts. However, each task varies in its composition of subprocesses and further each single cognitive subprocess may vary in its contribution to the observed static functional connectivity (i.e., connectivity averaged over the whole duration of the resting- or task-related scan). How specific cognitive subprocesses may contribute to the observed relation between less brain network reconfiguration and higher intelligence or more specific cognitive abilities cannot be investigated by relating these general task concepts to a general factor of intelligence (Salthouse et al. 2015) but constitutes an interesting subject for further research. We therefore recommend future studies a) to include more task diversity, b) to include different difficulty levels of tasks, and c) to identify separate cognitive subprocesses within tasks to explore the impact these subprocesses may have on the association of brain network reconfiguration and intelligence (or other cognitive measures). Lastly, the analyses reported here might also be adapted to time-resolved connectivity and the analyses of momentary switches between cognitive states (Shine et al. 2019; Greene et al. 2020).

In sum, our study suggests that greater efficiency in the reconfiguration of functional brain networks in response to various external demands is associated with an increased capacity for cognition and intellectual performance. In general, superior performance may profit from fast and efficient neural processing. The here observed association between general intelligence and less task-induced brain network reconfiguration that holds across a broad variety of different cognitive demands can be interpreted as support for the assumption that the intrinsic brain network architecture of more intelligent people is per se closer to a network configuration as required by various external demands. We conclude that such a network architecture constitutes an optimal foundation for fast and efficient cognitive processing that ultimately contributes to intelligent behavior. Finally, the involvement of multiple brain systems suggests intelligence as an emergent whole-brain phenomenon. Taken together, our study proposes multi-task brain network reconfiguration as promising marker to further understand the mechanisms underlying human cognition.

## Acknowledgements

The authors thank the Human Connectome Project (Van Essen et al. 2013), WU-Minn Consortium (Principal Investigators: David Van Essen and Kamil Ugurbil; 1U54MH091657) funded by the 16 NIH Institutes and Centers that support the NIH Blueprint for Neuroscience Research; and by the McDonnell Center for Systems Neuroscience at Washington University, for providing data of the main sample, and all contributors to The Amsterdam Open MRI Collection (Snoek et al. 2021, Principal Investigator: H. Steven Scholte) for providing data of the replication samples.

## Author Contributions

J.T., J.F, O.S. and K.H. conceived of the study. J.T. analyzed the data. J.F. and O.S. preprocessed the data and provided theoretical input during analyses. J.T. and K.H. wrote the manuscript. K.H. developed the initial idea and acquired funding for the project. All authors were involved in the interpretation of results and provided feedback during manuscript preparation.

## References

Assem M, Blank IA, Mineroff Z, Ademoğlu A, Fedorenko E. 2020. Activity in the fronto-parietal multiple-demand network is robustly associated with individual differences in working memory and fluid intelligence. Cortex. 131:1–16.

Avery EW, Yoo K, Rosenberg MD, Greene AS, Gao S, Na DL, Scheinost D, Constable TR, Chun MM. 2020. Distributed Patterns of Functional Connectivity Predict Working Memory Performance in Novel Healthy and Memory-impaired Individuals. J Cogn Neurosci. 32:241–255.

Barch DM, Burgess GC, Harms MP, Petersen SE, Schlaggar BL, Corbetta M, Glasser MF, Curtiss S, Dixit S, Feldt C, et al. 2013. Function in the human connectome: Task-fMRI and individual differences in behavior. Neuroimage. 80:169–189.

Basten U, Hilger K, Fiebach CJ. 2015. Where smart brains are different: A quantitative meta-analysis of functional and structural brain imaging studies on intelligence. Intelligence. 51:10–27.

Basten U, Stelzel C, Fiebach CJ. 2013. Intelligence is differentially related to neural effort in the task-positive and the task-negative brain network. Intelligence. 41:517–528.

Biswal B, Zerrin Yetkin F, Haughton VM, Hyde JS. 1995. Functional connectivity in the motor cortex of resting human brain using echo-planar mri. Magn Reson Med. 34:537–541.

Blokland GAM, McMahon KL, Hoffman J, Zhu G, Meredith M, Martin NG, Thompson PM, de Zubicaray GI, Wright MJ. 2008. Quantifying the heritability of task-related brain activation and performance during the N-back working memory task: A twin fMRI study. Biol Psychol. 79:70–79.

Braun U, Schäfer A, Walter H, Erk S, Romanczuk-Seiferth N, Haddad L, Schweiger JI, Grimm O, Heinz A, Tost H, et al. 2015. Dynamic reconfiguration of frontal brain networks during executive cognition in humans. Proc Natl Acad Sci. 112:11678–11683.

Catani M, Dell’Acqua F, Thiebaut de Schotten M. 2013. A revised limbic system model for memory, emotion and behaviour. Neurosci Biobehav Rev. 37:1724–1737.

Cohen JR, D’Esposito M. 2016. The Segregation and Integration of Distinct Brain Networks and Their Relationship to Cognition. J Neurosci. 36:12083–12094.

Cole MW, Bassett DS, Power JD, Braver TS, Petersen SE. 2014. Intrinsic and Task-Evoked Network Architectures of the Human Brain. Neuron. 83:238–251.

Cole MW, Ito T, Cocuzza C, Sanchez-Romero R. 2021. The Functional Relevance of Task-State Functional Connectivity. J Neurosci. 41:2684–2702.

Cole MW, Ito T, Schultz D, Mill R, Chen R, Cocuzza C. 2019. Task activations produce spurious but systematic inflation of task functional connectivity estimates. Neuroimage. 189:1–18.

Dubois J, Galdi P, Paul LK, Adolphs R. 2018. A distributed brain network predicts general intelligence from resting-state human neuroimaging data. Philos Trans R Soc B Biol Sci. 373:20170284.

Duncan J. 2010. The multiple-demand (MD) system of the primate brain: mental programs for intelligent behaviour. Trends Cogn Sci. 14:172–179.

Dunst B, Benedek M, Jauk E, Bergner S, Koschutnig K, Sommer M, Ischebeck A, Spinath B, Arendasy M, Bühner M, et al. 2014. Neural efficiency as a function of task demands. Intelligence. 42:22–30.

Esteban O, Markiewicz CJ, Blair RW, Moodie CA, Isik AI, Erramuzpe A, Kent JD, Goncalves M, DuPre E, Snyder M, et al. 2019. fMRIPrep: a robust preprocessing pipeline for functional MRI. Nat Methods. 16:111–116.

Finn ES, Shen X, Scheinost D, Rosenberg MD, Huang J, Chun MM, Papademetris X, Constable RT. 2015. Functional connectome fingerprinting: identifying individuals using patterns of brain connectivity. Nat Neurosci. 18:1664–1671.

Gao S, Greene AS, Constable RT, Scheinost D. 2019. Combining multiple connectomes improves predictive modeling of phenotypic measures. Neuroimage. 201:116038.

Glasser MF, Sotiropoulos SN, Wilson JA, Coalson TS, Fischl B, Andersson JL, Xu J, Jbabdi S, Webster M, Polimeni JR, et al. 2013. The minimal preprocessing pipelines for the Human Connectome Project. Neuroimage. 80:105–124.

Greene AS, Gao S, Noble S, Scheinost D, Constable RT. 2020. How Tasks Change Whole-Brain Functional Organization to Reveal Brain-Phenotype Relationships. Cell Rep. 32:108066.

Greene AS, Gao S, Scheinost D, Constable RT. 2018. Task-induced brain state manipulation improves prediction of individual traits. Nat Commun. 9:2807.

Gregory MD, Kippenhan JS, Dickinson D, Carrasco J, Mattay VS, Weinberger DR, Berman KF. 2016. Regional Variations in Brain Gyrification Are Associated with General Cognitive Ability in Humans. Curr Biol. 26:1301–1305.

Haier RJ. 2017. The Neuroscience of Intelligence. Cambridge (UK): Cambridge University Press.

Haier RJ, Siegel B V., Nuechterlein KH, Hazlett E, Wu JC, Paek J, Browning HL, Buchsbaum MS. 1988. Cortical glucose metabolic rate correlates of abstract reasoning and attention studied with positron emission tomography. Intelligence. 12:199–217.

Hearne LJ, Cocchi L, Zalesky A, Mattingley JB. 2017. Reconfiguration of Brain Network Architectures between Resting-State and Complexity-Dependent Cognitive Reasoning. J Neurosci. 37:8399–8411.

Hilger K, Ekman M, Fiebach CJ, Basten U. 2017a. Intelligence is associated with the modular structure of intrinsic brain networks. Sci Rep. 7:16088.

Hilger K, Ekman M, Fiebach CJ, Basten U. 2017b. Efficient hubs in the intelligent brain: Nodal efficiency of hub regions in the salience network is associated with general intelligence. Intelligence. 60:10–25.

Hilger K, Fukushima M, Sporns O, Fiebach CJ. 2020. Temporal stability of functional brain modules associated with human intelligence. Hum Brain Mapp. 41:362–372.

Hu L, Bentler PM. 1999. Cutoff criteria for fit indexes in covariance structure analysis: Conventional criteria versus new alternatives. Struct Equ Model A Multidiscip J. 6:1–55.

Jenkinson M, Bannister P, Brady M, Smith S. 2002. Improved Optimization for the Robust and Accurate Linear Registration and Motion Correction of Brain Images. Neuroimage. 17:825–841.

Jensen AR. 2006. Clocking the Mind. Elsevier.

Jung RE, Haier RJ. 2007. The Parieto-Frontal Integration Theory (P-FIT) of intelligence: Converging neuroimaging evidence. Behav Brain Sci. 30:135–154.

Kovacs K, Conway ARA. 2016. Process Overlap Theory: A Unified Account of the General Factor of Intelligence. Psychol Inq. 27:151–177.

Liu T, Pleskac TJ. 2011. Neural correlates of evidence accumulation in a perceptual decision task. J Neurophysiol. 106:2383–2398.

Lustig C, Buckner RL. 2004. Preserved Neural Correlates of Priming in Old Age and Dementia. Neuron. 42:865–875.

Marek S, Tervo-Clemmens B, Calabro FJ, Montez DF, Kay BP, Hatoum AS, Donohue MR, Foran W, Miller RL, Feczko E, et al. 2020. Towards Reproducible Brain-Wide Association Studies. bioRxiv.

Neubauer AC, Fink A. 2009. Intelligence and neural efficiency: Measures of brain activation versus measures of functional connectivity in the brain. Intelligence. 37:223–229.

Parkes L, Fulcher B, Yücel M, Fornito A. 2018. An evaluation of the efficacy, reliability, and sensitivity of motion correction strategies for resting-state functional MRI. Neuroimage. 171:415–436.

Ragland JD, Turetsky BI, Gur RC, Gunning-Dixon F, Turner T, Schroeder L, Chan R, Gur RE. 2002. Working memory for complex figures: An fMRI comparison of letter and fractal n-back tasks. Neuropsychology. 16:370–379.

Raven JC, Court JH. 1998. Manual for Raven’s progressive matrices and vocabulary scales.

Salehi M, Karbasi A, Barron DS, Scheinost D, Constable RT. 2020. Individualized functional networks reconfigure with cognitive state. Neuroimage. 206:116233.

Salthouse TA, Habeck C, Razlighi Q, Barulli D, Gazes Y, Stern Y. 2015. Breadth and age-dependency of relations between cortical thickness and cognition. Neurobiol Aging. 36:3020–3028.

Santarnecchi E, Emmendorfer A, Pascual-Leone A. 2017. Dissecting the parieto-frontal correlates of fluid intelligence: A comprehensive ALE meta-analysis study. Intelligence. 63:9–28.

Santarnecchi E, Momi D, Mencarelli L, Plessow F, Saxena S, Rossi S, Rossi A, Mathan S, Pascual-Leone A. 2021. Overlapping and dissociable brain activations for fluid intelligence and executive functions. Cogn Affect Behav Neurosci.

Schaefer A, Kong R, Gordon EM, Laumann TO, Zuo X-N, Holmes AJ, Eickhoff SB, Yeo BTT. 2018. Local-Global Parcellation of the Human Cerebral Cortex from Intrinsic Functional Connectivity MRI. Cereb Cortex. 28:3095–3114.

Schmid J, Leiman JM. 1957. The development of hierarchical factor solutions. Psychometrika. 22:53–61.

Schubert A-L, Frischkorn GT. 2020. Neurocognitive Psychometrics of Intelligence: How Measurement Advancements Unveiled the Role of Mental Speed in Intelligence Differences. Curr Dir Psychol Sci. 29:140–146.

Schultz DH, Cole MW. 2016. Higher intelligence is associated with less task-related brain network reconfiguration. J Neurosci. 36:8551–8561.

Shen X, Finn ES, Scheinost D, Rosenberg MD, Chun MM, Papademetris X, Constable RT. 2017. Using connectome-based predictive modeling to predict individual behavior from brain connectivity. Nat Protoc. 12:506–518.

Shine JM, Bissett PG, Bell PT, Koyejo O, Balsters JH, Gorgolewski KJ, Moodie CA, Poldrack RA. 2016. The Dynamics of Functional Brain Networks: Integrated Network States during Cognitive Task Performance. Neuron. 92:544–554.

Shine JM, Breakspear M, Bell PT, Ehgoetz Martens KA, Shine R, Koyejo O, Sporns O, Poldrack RA. 2019. Human cognition involves the dynamic integration of neural activity and neuromodulatory systems. Nat Neurosci. 22:289–296.

Smith SM, Beckmann CF, Andersson J, Auerbach EJ, Bijsterbosch J, Douaud G, Duff E, Feinberg DA, Griffanti L, Harms MP, et al. 2013. Resting-state fMRI in the Human Connectome Project. Neuroimage. 80:144–168.

Snoek L, van der Miesen MM, Beemsterboer T, van der Leij A, Eigenhuis A, Steven Scholte H. 2021. The Amsterdam Open MRI Collection, a set of multimodal MRI datasets for individual difference analyses. Sci Data. 8:85.

Soreq E, Violante IR, Daws RE, Hampshire A. 2021. Neuroimaging evidence for a network sampling theory of individual differences in human intelligence test performance. Nat Commun. 12:2072.

Spearman C. 1904. “General Intelligence,” Objectively Determined and Measured. Am J Psychol. 15:201.

Sripada C, Angstadt M, Rutherford S, Taxali A, Shedden K. 2020. Toward a “treadmill test” for cognition: Improved prediction of general cognitive ability from the task activated brain. Hum Brain Mapp. 41:3186–3197.

Sternberg RJ. 1997. The concept of intelligence and its role in lifelong learning and success. Am Psychol. 52:1030–1037.

Unsworth N, Miller JD, Lakey CE, Young DL, Meeks JT, Campbell WK, Goodie AS. 2009. Exploring the Relations Among Executive Functions, Fluid Intelligence, and Personality. J Individ Differ. 30:194–200.

Van den Heuvel MP, Hulshoff Pol HE. 2010. Exploring the brain network: A review on resting-state fMRI functional connectivity. Eur Neuropsychopharmacol. 20:519–534.

Van Essen DC, Smith SM, Barch DM, Behrens TEJ, Yacoub E, Ugurbil K. 2013. The WU-Minn Human Connectome Project: An overview. Neuroimage. 80:62–79.

Yeo TBT, Krienen FM, Sepulcre J, Sabuncu MR, Lashkari D, Hollinshead M, Roffman JL, Smoller JW, Zöllei L, Polimeni JR, et al. 2011. The organization of the human cerebral cortex estimated by intrinsic functional connectivity. J Neurophysiol. 106:1125–1165.

