## Supplementary Figures and Tables for "Multi-Task Brain Network Reconfiguration is Inversely Associated with Human Intelligence"

Running title: Brain Network Reconfiguration and Human Intelligence


Jonas A. Thiele[1]*, Joshua Faskowitz[2], Olaf Sporns[2], Kirsten Hilger[1]*

[1] Department of Psychology I - Biological Psychology, Clinical Psychology and Psychotherapy, Marcusstr. 9-11, 97070 Würzburg, Germany

[2] Department of Psychological and Brain Sciences, Indiana University, Psychology Building 360, 1101 E 10th Street, Bloomington, IN 47405, USA

**ORCID:**

Jonas Thiele: 0000-0003-2702-9690

Joshua Faskowitz: 0000-0003-1814-7206

Olaf Sporns: 0000-0001-7265-4036

Kirsten Hilger: 0000-0003-3940-5884

* Corresponding authors:

Jonas Alexander Thiele & Kirsten Hilger

Department of Psychology I

Marcusstr. 9-11

D-97070 Würzburg

**Table S1.** Cognitive measures from the Human Connectome Project (HCP, Barch et al. 2013) used for estimating a latent factor of general intelligence (*g*-factor).

| Test | Instrument | Measure used |
|---|---|---|
| **1** | Episodic Memory (Picture Sequence Memory) | PicSeq_Unadj |
| **2** | Executive Function/Cognitive Flexibility (Dimensional Change Card Sort) | CardSort_Unadj |
| **3** | Executive Function/Inhibition (Flanker Task) | Flanker_Unadj |
| **4** | Fluid Intelligence (Penn Progressive Matrices) | PMAT24_A_CR |
| **5** | Language/Reading Decoding (Oral Reading Recognition) | ReadEng_Unadj |
| **6** | Language/Vocabulary Comprehension (Picture Vocabulary) | PicVocab_Unadj |
| **7** | Processing Speed (Pattern Completion Processing Speed) | ProcSpeed_Unadj |
| **8** | Self-regulation/Impulsivity (Delay Discounting) | DDisc_AUC_200 + DDisc_AUC_40K |
| **9** | Spatial Orientation (Variable Short Penn Line Orientation Test) | VSPLOT_TC |
| **10** | Sustained Attention (Short Penn Continuous Performance Test) | $\frac{\mathrm{SCPT\_TP} + \mathrm{SCPT\_TN}}{(\mathrm{SCPT\_TP} + \mathrm{SCPT\_TN} + \mathrm{SCPT\_FP} + \mathrm{SCPT\_FN})\mathrm{SCPT\_TPRT}}$ |
| **11** | Verbal Episodic Memory (Penn Word Memory Test) | IWRD_TOT |
| **12** | Working Memory (List Sorting) | ListSort_Unadj |

**Table S2.** Overview over main-, control-, and replication-analyses results, consistently demonstrating that less brain network reconfiguration is associated with higher intelligence scores.

| **Sample** | **FC filtering procedure** | ***rho* (*p*) rest-task reconfiguration** | ***rho* (*p*) task-task reconfiguration** |
|---|---|---|---|
| **Main sample** – main analysis | correlation with intelligence *p* < .1 | -.23 (< .001) | -.23 (< .001) |
| **Main sample** – control analyses | correlation with intelligence *p* < .01 | -.19 (< .001) | -.23 (< .001) |
| | correlation with intelligence *p* < .1, no removal of instable connections | -.21 (< .001) | -.21 (< .001) |
| | no filter | -.12 (< .001) | -.12 (< .001) |
| **Replication sample 1** (PIOP1) | intersection with robust main sample mask | -.32 (< .001) | -.26 (< .001) |
| **Replication sample 2** (PIOP2) | intersection with robust main sample mask | -.23 (< .001) | -.26 (< .001) |

*Note.* Association (partial Spearman correlation, *rho*) between a latent *g*-factor of intelligence (derived from 12 tasks, see Table S1, Fig. S2) and brain network reconfiguration operationalized as cosine distance between functional connectivity matrices (FCs) averaged over all possible rest-task combinations, and all possible task-task combinations, respectively. All correlations were controlled for influences of age, sex, handedness, and in-scanner head motion (mean framewise displacement). Results from different filtering procedures, i.e., techniques for selecting relevant functional brain connections, and results from the replication analyses are shown. In the main analysis, all connections correlating with intelligence *p* < .1 were included, except connections that were inconsistently correlated with intelligence, i.e., in both positive and negative direction in different states. For additional control analysis, this FC filtering approach was repeated a) with a *p*-threshold of *p* < .01, b) with a *p*-threshold of *p* < .1 but without the removal of inconsistently correlated connections, and c) without any filtering. In the replication analyses no *p*-threshold was used, but instead a filter mask was computed from the larger main sample (containing connections correlating only either positively or negatively with intelligence *p* < .01 in at least one of the filtered FCs of intersecting state combinations) and only connections located in these main sample filter mask and correlating with intelligence in the same direction in the replication samples were used in further analyses.

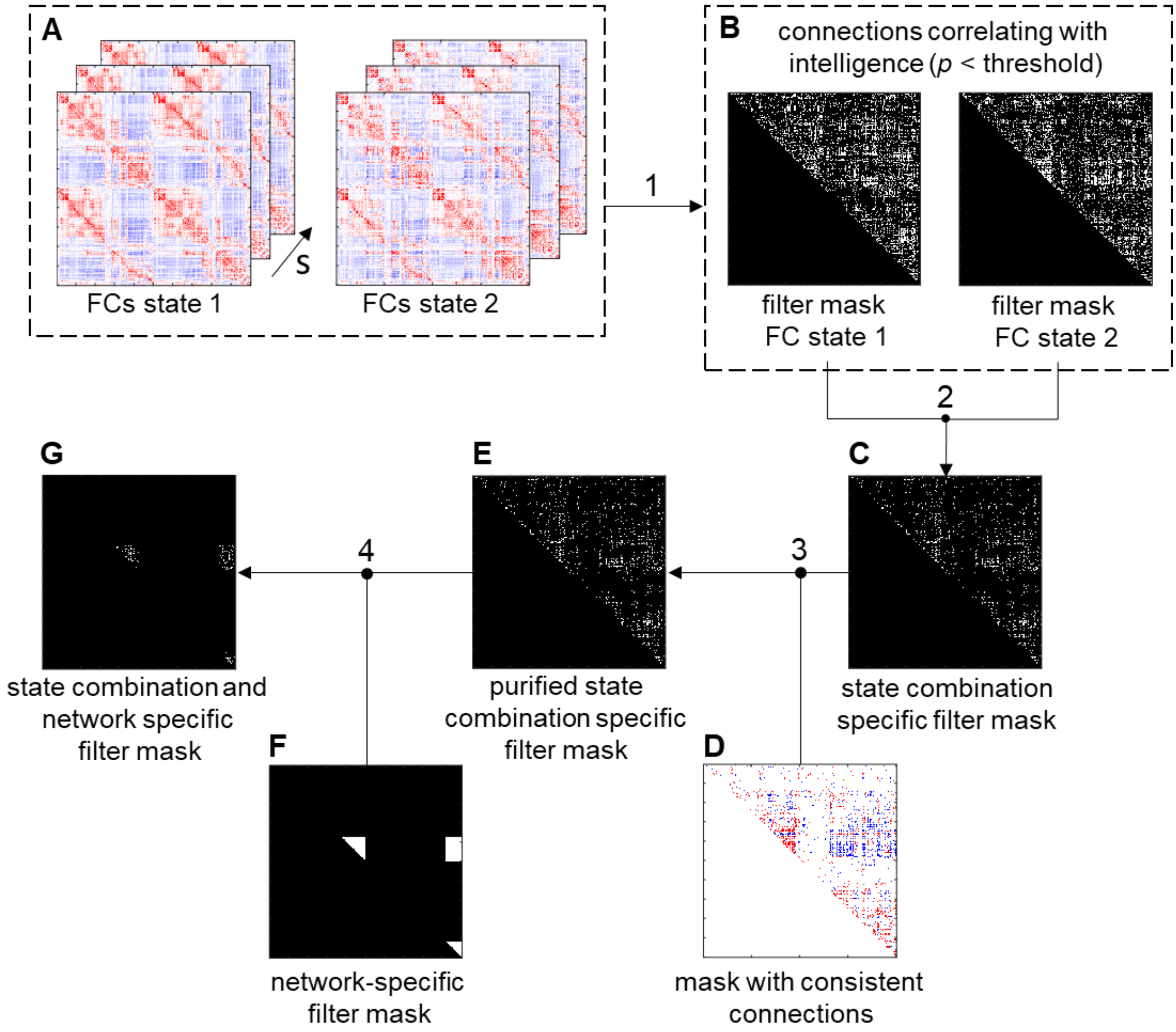


**Fig S1.** Schematic illustration of the functional connectivity filtering procedure. To prevent circularity in the process of selecting connections, the filtering was 10-fold cross-validated. For this reason, the sample was divided into ten subsamples. For each fold, filter masks were computed on nine subsamples and applied to the withheld sample, respectively. In each fold, the subject-specific functional connectivity matrices (FCs) of the two states that were compared with each other (resting state and one task, or two tasks, **A**) were correlated with intelligence (1) separately using data from nine sub-samples. Within each of these matrices, functional brain connections significantly correlated with intelligence ($p$ < threshold, see Methods) were kept resulting in two state-specific filter masks (**B**, remaining connections are displayed in white) for this specific fold. A state combination specific filter mask (**C**), as necessary for computation of state combination specific reconfiguration measures, was derived by intersecting the two state-specific filter masks (2) of this fold. Potential noise was then removed by excluding connections that were, in this fold, not consistently correlated with intelligence across states (i.e., in one state positive and in another state negative, 3). For this reason, a mask, containing connections correlating only either positively (red) or negatively (blue) with intelligence $p < .01$ in at least one of all states (rest and seven tasks) was used as reference (**D**). This resulted in the purified state combination specific filter mask (**E**). For computing whole-brain reconfiguration, the FCs of both compared states from the withheld sub-sample were filtered with this purified state combination specific filter mask (**E**). For computing network-specific brain reconfiguration measures, a state combination and network specific filter mask (**G**) was created by intersecting (4) the purified state combination specific filter mask (**E**) and a filter mask including all connections belonging to a specific functional brain network, i.e., within the network or between this network and another network (**F**) and applied to the withheld subsample.

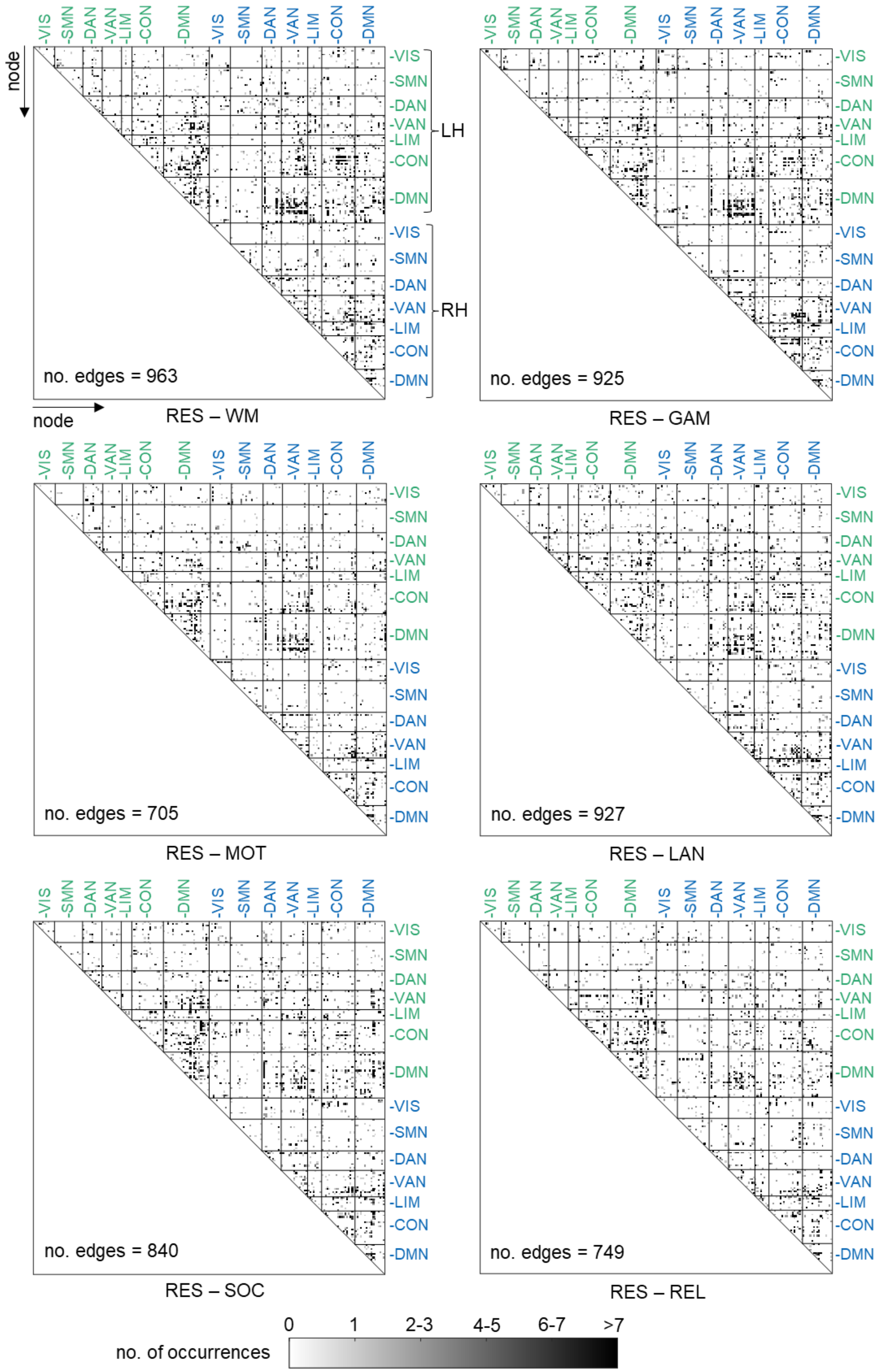
node
-VIS
-SMN
-DAN
-VAN
-LIM
-CON
-DMN
LH
RH
no. edges = 963
RES – WM
no. edges = 925
RES – GAM
no. edges = 705
RES – MOT
no. edges = 927
RES – LAN
no. edges = 840
RES – SOC
no. edges = 749
RES – REL
no. of occurrences
0
1
2-3
4-5
6-7
>7

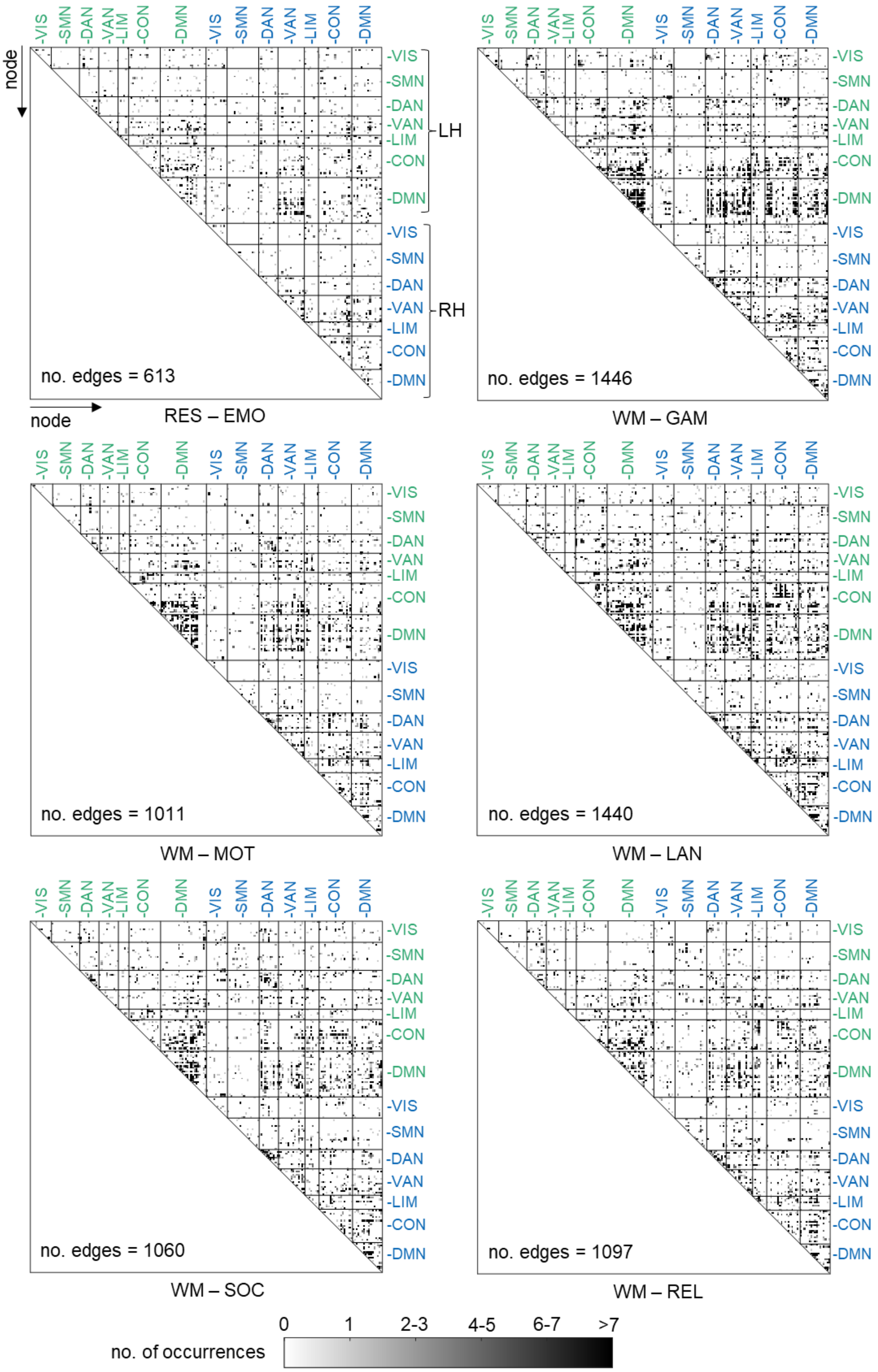

-VIS -SMN -DAN -VAN -LIM -CON -DMN -VIS -SMN -DAN -VAN -LIM -CON -DMN
LH
RH
node
node
no. edges = 613
RES – EMO
no. edges = 1446
WM – GAM
no. edges = 1011
WM – MOT
no. edges = 1440
WM – LAN
no. edges = 1060
WM – SOC
no. edges = 1097
WM – REL
no. of occurrences
0
1
2-3
4-5
6-7
>7

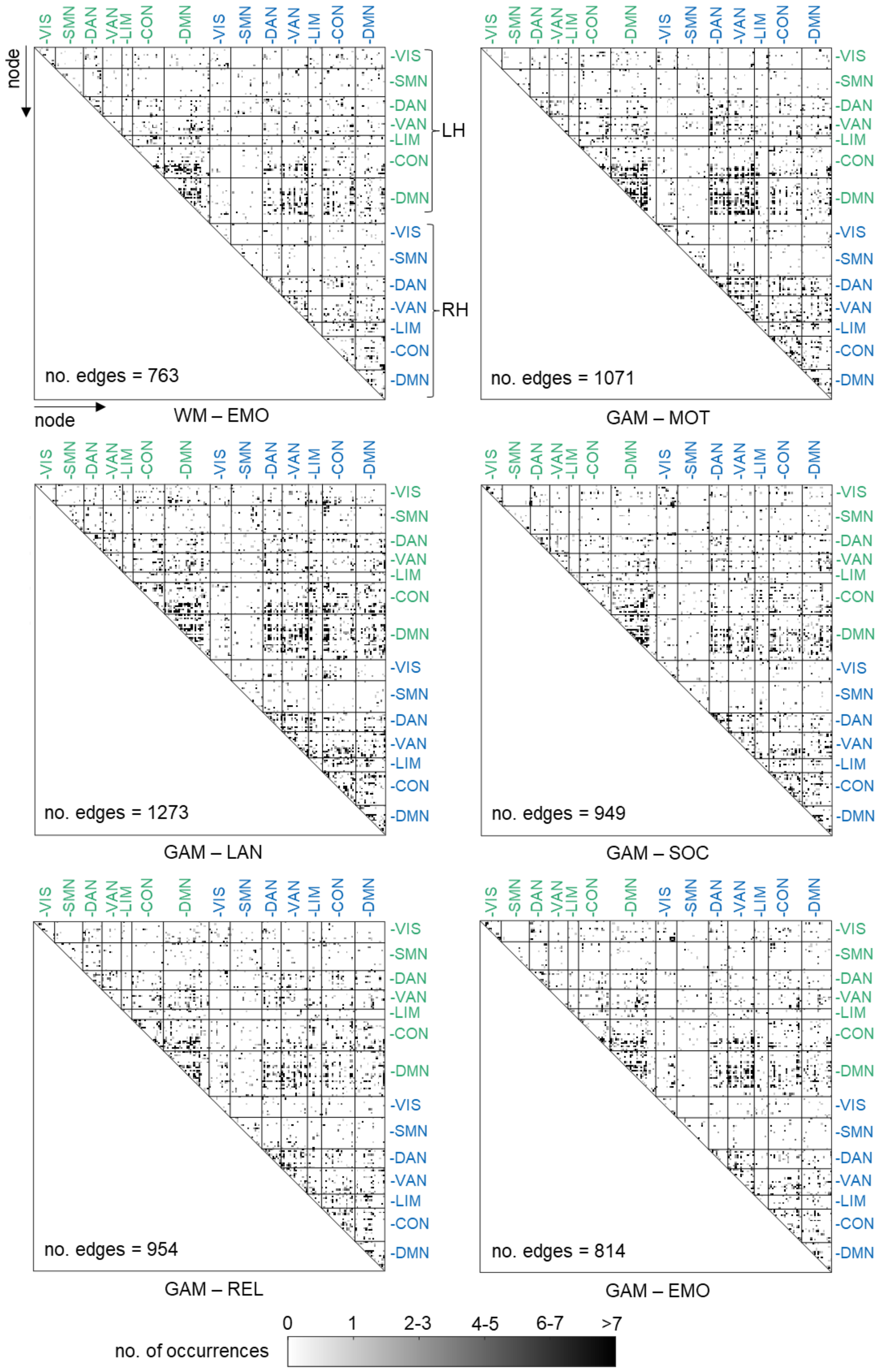
node
node
-VIS
-SMN
-DAN
-VAN
-LIM
-CON
-DMN
LH
RH
no. edges = 763
WM – EMO
no. edges = 1071
GAM – MOT
no. edges = 1273
GAM – LAN
no. edges = 949
GAM – SOC
no. edges = 954
GAM – REL
no. edges = 814
GAM – EMO
no. of occurrences
0
1
2-3
4-5
6-7
>7

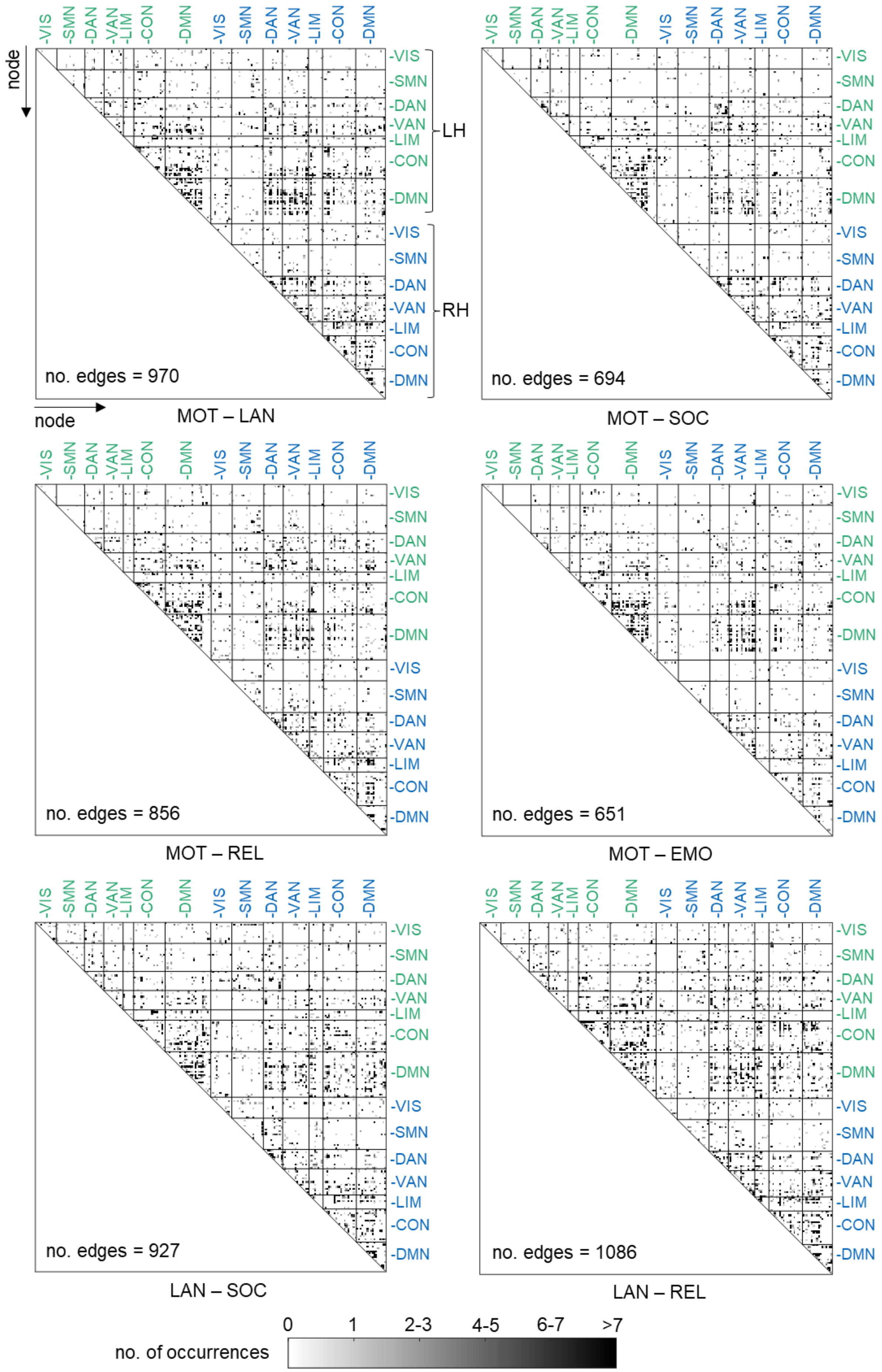
node
VIS
SMN
DAN
VAN
LIM
CON
DMN
LH
RH
no. edges = 970
MOT – LAN
no. edges = 694
MOT – SOC
no. edges = 856
MOT – REL
no. edges = 651
MOT – EMO
no. edges = 927
LAN – SOC
no. edges = 1086
LAN – REL
no. of occurrences
0
1
2-3
4-5
6-7
>7

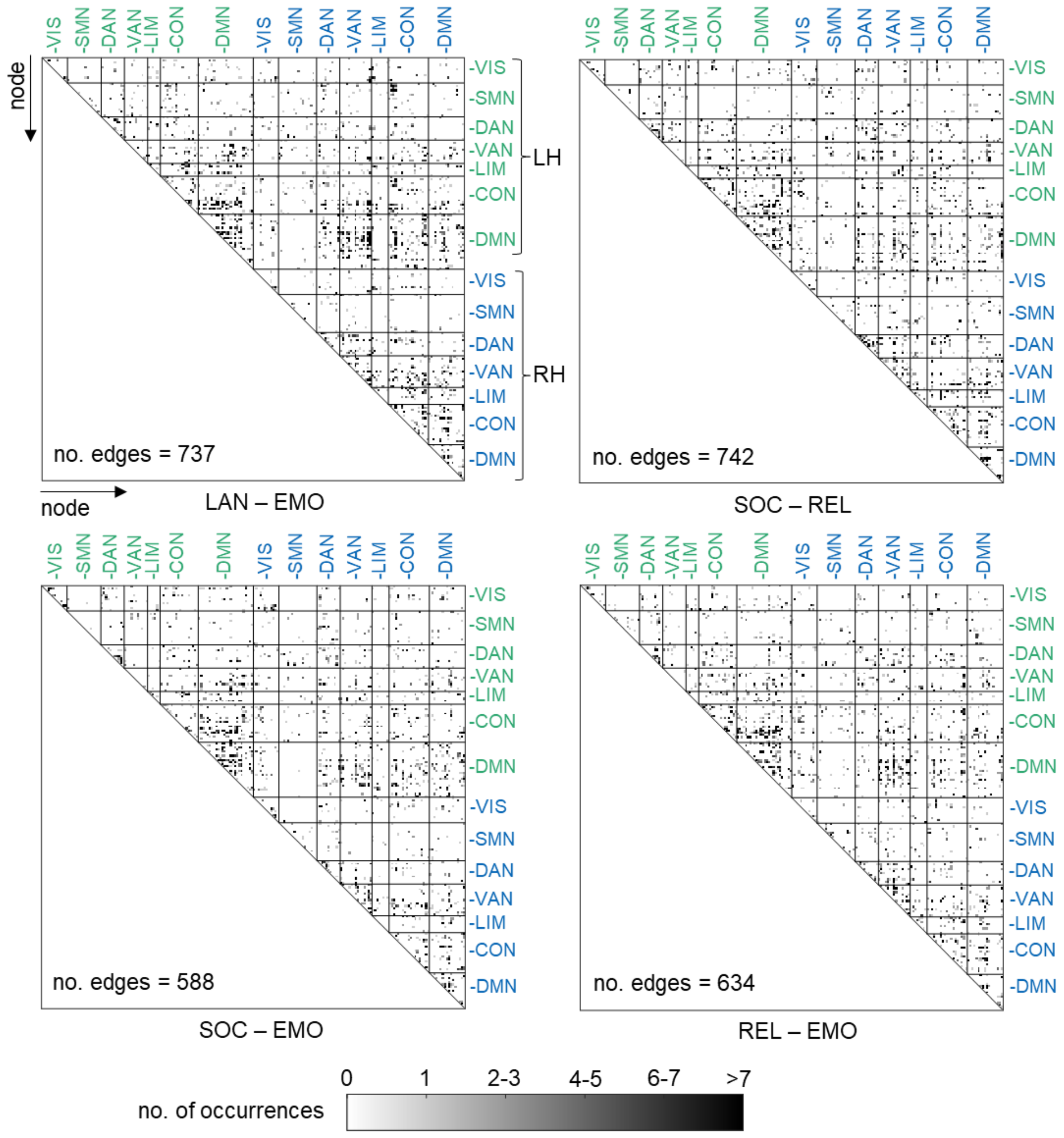


**Fig. S2.** Remaining and filtered functional brain connections for each state combination averaged over all ten cross-validation folds. Functional connections were filtered based on their correlation with intelligence ($p$ < .1, Finn et al. 2015). Connections inconsistently correlated with intelligence across states (positive in one, negative in another or vice versa) were excluded. Connections between regions that remain after filtering are saturated according to their number of occurrences across all folds. Corresponding networks are displayed on the x- and y-axes (seven network partition, Yeo et al. 2011). Black lines indicate the boarders between different networks. The number of remaining connections (averaged over ten folds) is depicted in the lower left corner of each subfigure. Abbreviations of the two cognitive states of the respective state combination are displayed below each subfigure. VIS, visual network; SMN, somatomotor network; DAN, dorsal attention network; VAN, salience/ventral attention network; LIM, limbic network; CON, control network; DMN, default mode network; RES, resting state; WM, working memory task; GAM, gambling task; MOT, motor task; LAN, language processing task; SOC, social cognition task; REL, relational processing task; EMO, emotion processing task; LH, right hemisphere; LH, left hemisphere.

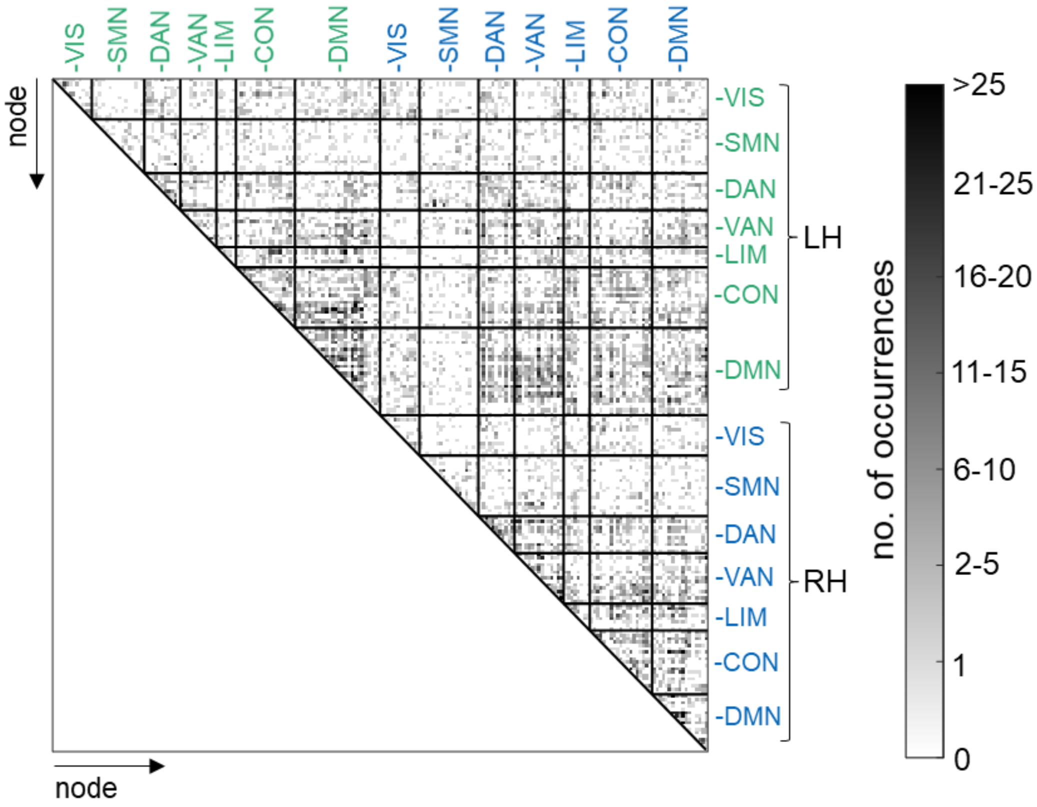


**Fig. S3.** Remaining functional brain connections summed over all state combinations. Functional connections were filtered based on their correlation with intelligence (*p* < .1, Finn et al. 2015). Connections inconsistently correlated with intelligence across states (positive in one, negative in another or vice versa) were excluded. Corresponding networks are displayed on the x- and y-axes (seven network partition, Yeo et al. 2011). Black lines indicate the boarders between different networks. Connections between regions that remain after filtering are saturated according to their number of occurrences across all state combinations (averaged over all ten cross-validation folds of the filtering procedure, respectively). VIS, visual network; SMN, somatomotor network; DAN, dorsal attention network; VAN, salience/ventral attention network; LIM, limbic network; CON, control network; DMN, default mode network; LH, right hemisphere; LH, left hemisphere.

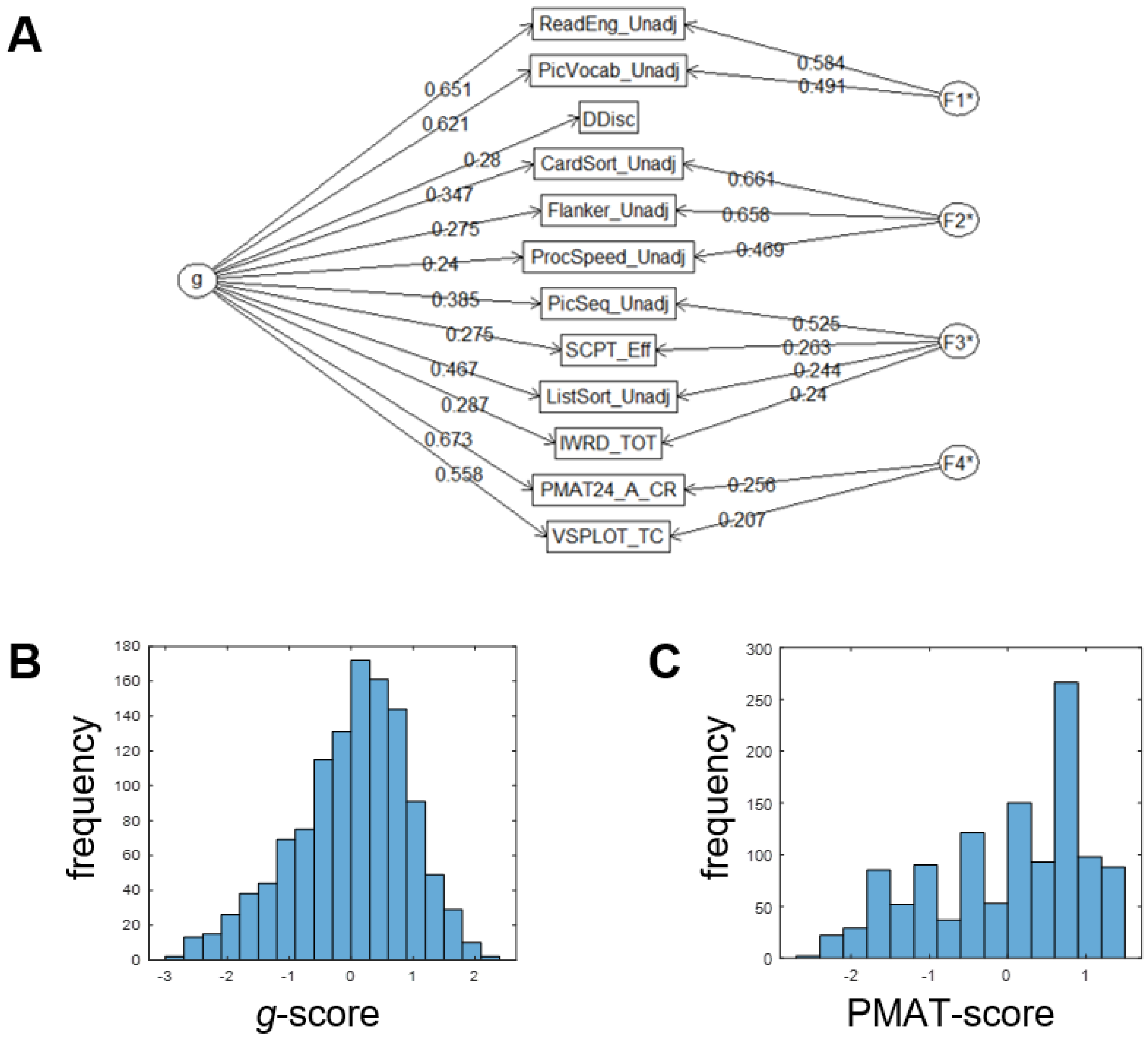


**Fig. S4.** Estimation of general intelligence (*g*-factor) from 12 cognitive scores (see Table S1). (**A**) Bi-factor model. (**B**) Distribution of estimated *g*-scores of 1186 subjects. (**C**) Distribution of PMAT-scores (Penn Progressive Matrices, brief assessment of intelligence provided by the HCP, Barch et al. 2013) of the sample.

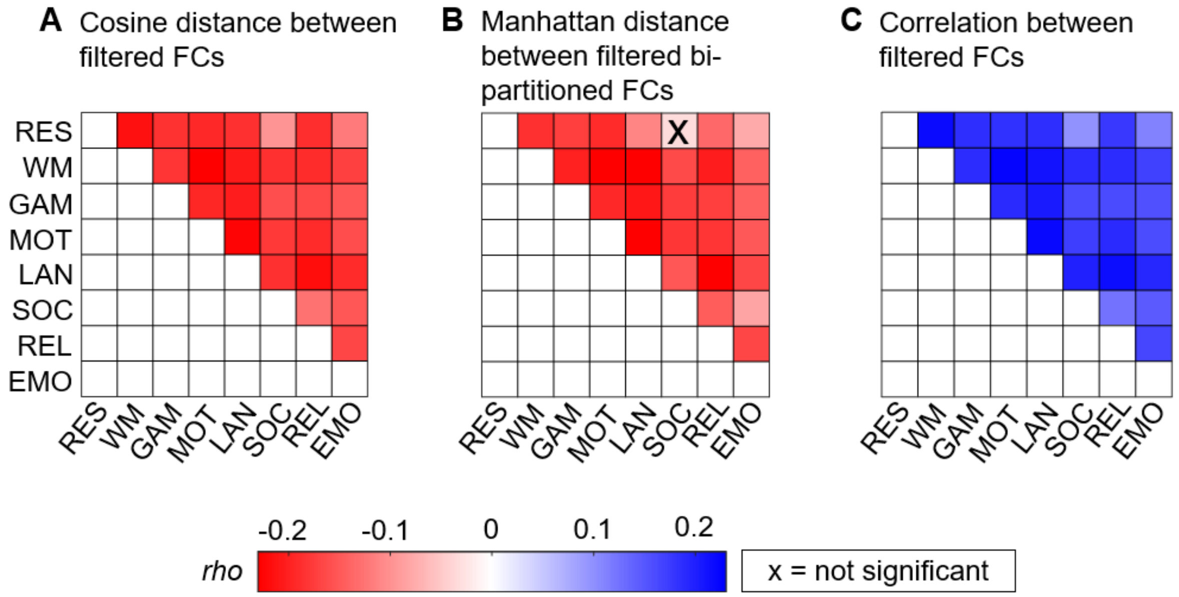


**Fig. S5.** Intelligence and state-specific brain network reconfiguration for different reconfiguration measures. Spearman correlation between intelligence and brain network reconfiguration for all possible state combinations (FDR-corrected *p*-values, *α* = .05). The strengths of correlations are depicted in different colors (see color bar). All correlations were controlled for influences of age, sex, handedness, and in-scanner head motion (mean framewise displacement) and reached significance except the single marked correlation. Reconfiguration was operationalized as (**A**) the cosine distance between filtered functional connectivity (FC) of state combinations, (**B**) the Manhattan distance between filtered bi-partitioned FCs of state combinations, and (**C**) as the Pearson correlation between filtered FCs of state combinations. FDR, false discovery rate; RES, resting state; WM, working memory task; GAM, gambling task; MOT, motor task; LAN, language processing task; SOC, social cognition task; REL, relational processing task; EMO, emotion processing task.

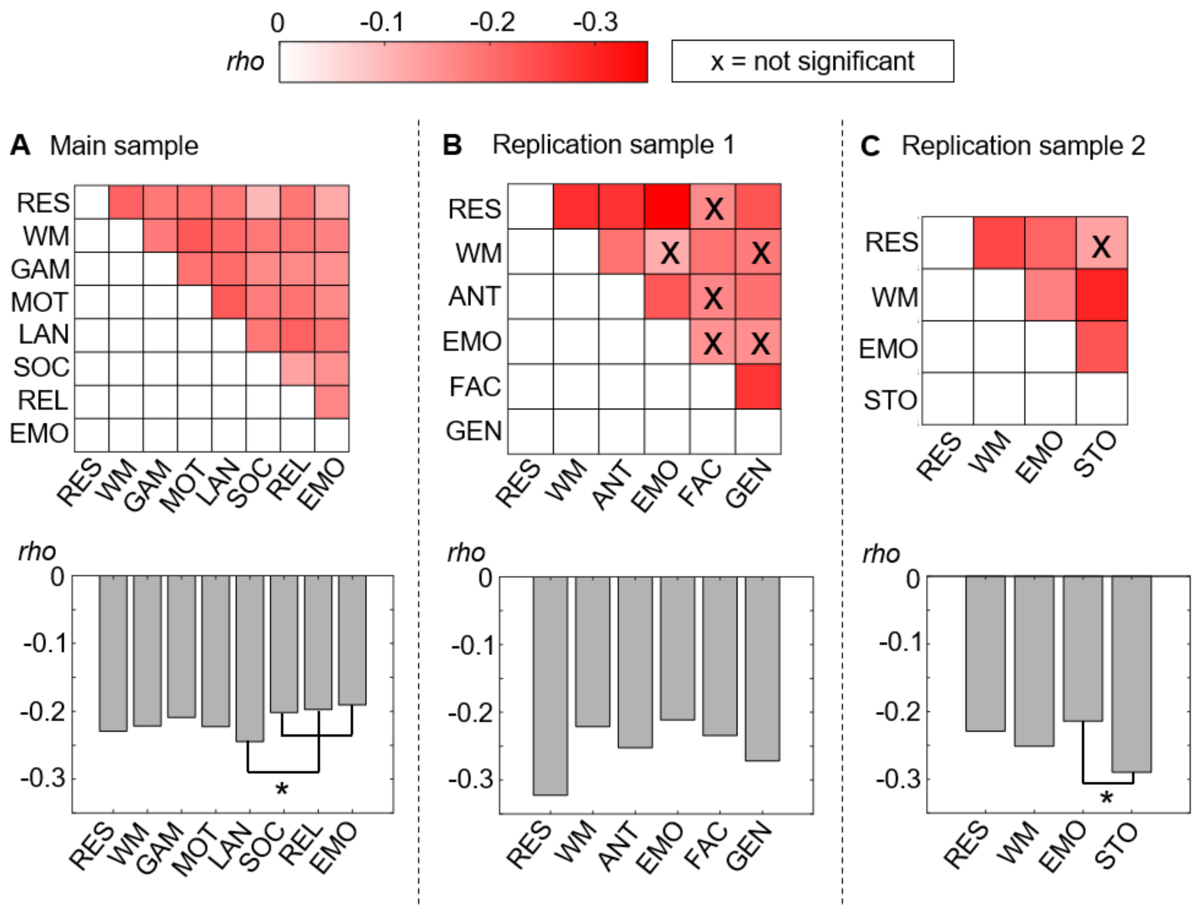


**Fig. S6.** Intelligence and state-specific reconfiguration across main and replication samples. Results are shown for (**A**) the main sample (similar to Fig. 2B), (**B**) the replication sample 1 (PIOP1), and (**C**) the replication sample 2 (PIOP2). Upper panel: Correlation between intelligence and brain network reconfiguration for all possible state combinations (FDR-corrected *p*-values, $\alpha = .05$). The strengths of correlations are depicted in different colors (see color bar). All correlations were controlled for influences of age, sex, handedness, and in-scanner head motion (mean framewise displacement) and reached significance except correlations marked with a black cross. Lower panel: Correlations between intelligence and a total measure of state-specific reconfiguration (FDR-corrected *p*-values, $\alpha = .05$, all correlations are significant). Specifically, reconfiguration values were averaged over all state combinations the respective state was involved in. Note that for task states, only combinations with different tasks (no rest) were included. Significant differences in correlation values ($p < .05$, marked with an asterisk) were only observed in the main sample for the comparisons between the language task (LAN) and the social cognition, the relational processing, and the emotion processing task (SOC, REL, EMO) respectively and in the replication sample 2 between the emotion task and the stop signal task. FDR, false discovery rate; RES, resting state; WM, working memory task; GAM, gambling task; MOT, motor task; LAN, language processing task; SOC, social cognition task; REL, relational processing task; EMO, emotion task; ANT, anticipation task; FAC, face perception task; GEN, gender-stroop task; STO, stop signal task.

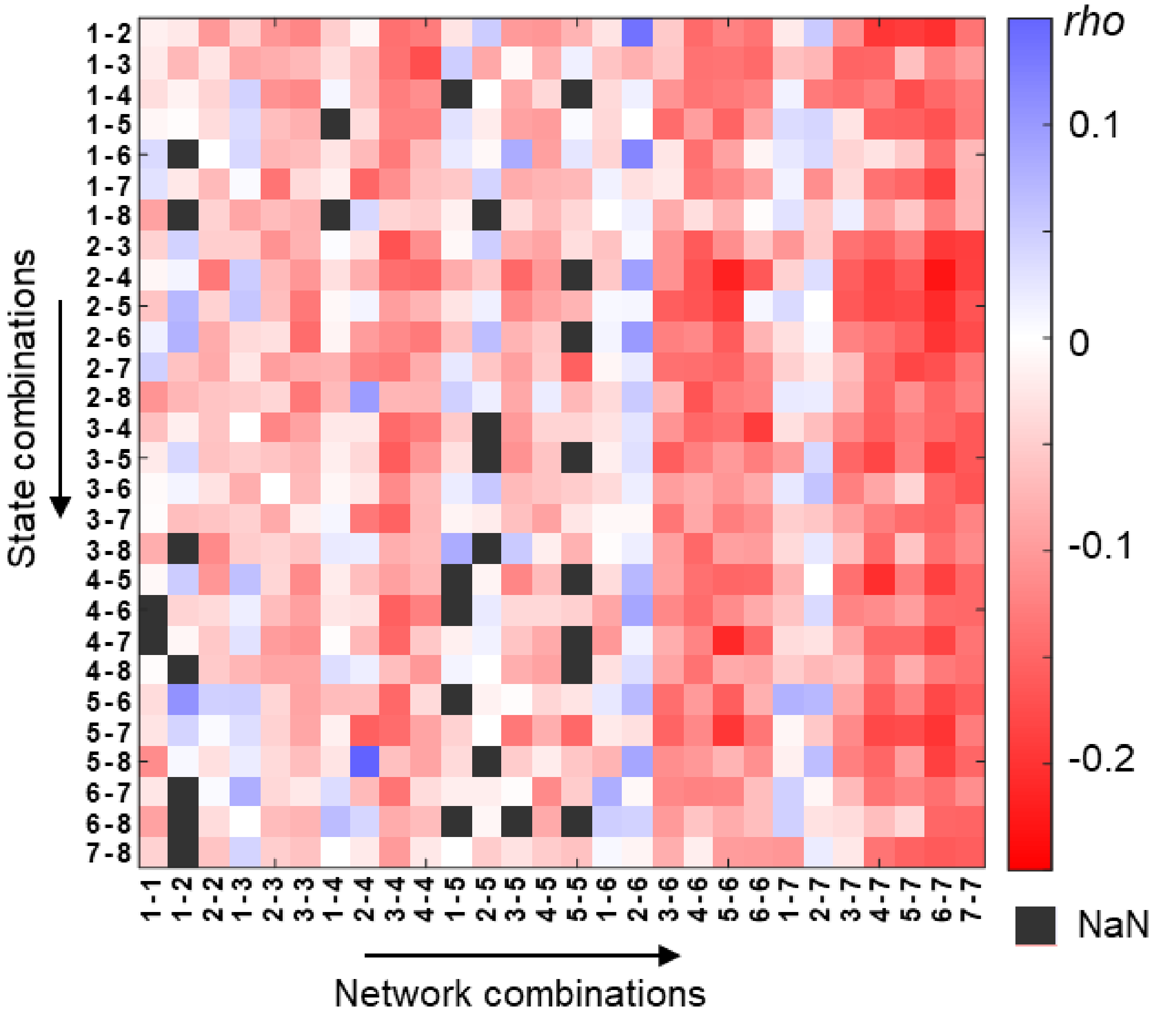


**Fig. S7.** Intelligence and brain network- and state-specific reconfiguration values. Note that this figure is similar to Fig. 2C but provides a more detailed description of the respective state and network combinations. Brain networks were derived from the Yeo atlas (Yeo et al. 2011, seven network partition used here) and network combinations refer to all within and between network connectivity combinations (columns). The strengths of the associations (Spearman correlations, *rho*) are depicted in different colors (see color bar). All correlations were controlled for influences of age, sex, handedness, and in-scanner head motion (mean framewise displacement). Note that NaN (not a number) values exist if in a specific network-state combination no single brain connection passes the filtering procedure (see Methods).The assignment of the numbers on the plot's axes to states or networks is the following: States, 1: resting state; 2: working memory task; 3: gambling task; 4: motor task; 5: language processing task; 6: social cognition task; 7: relational processing task; 8: emotion processing task. Networks, 1: visual network; 2: somatomotor network; 3: dorsal attention network; 4: salience/ventral attention network; 5: limbic network; 6: control network; 7: default mode network.

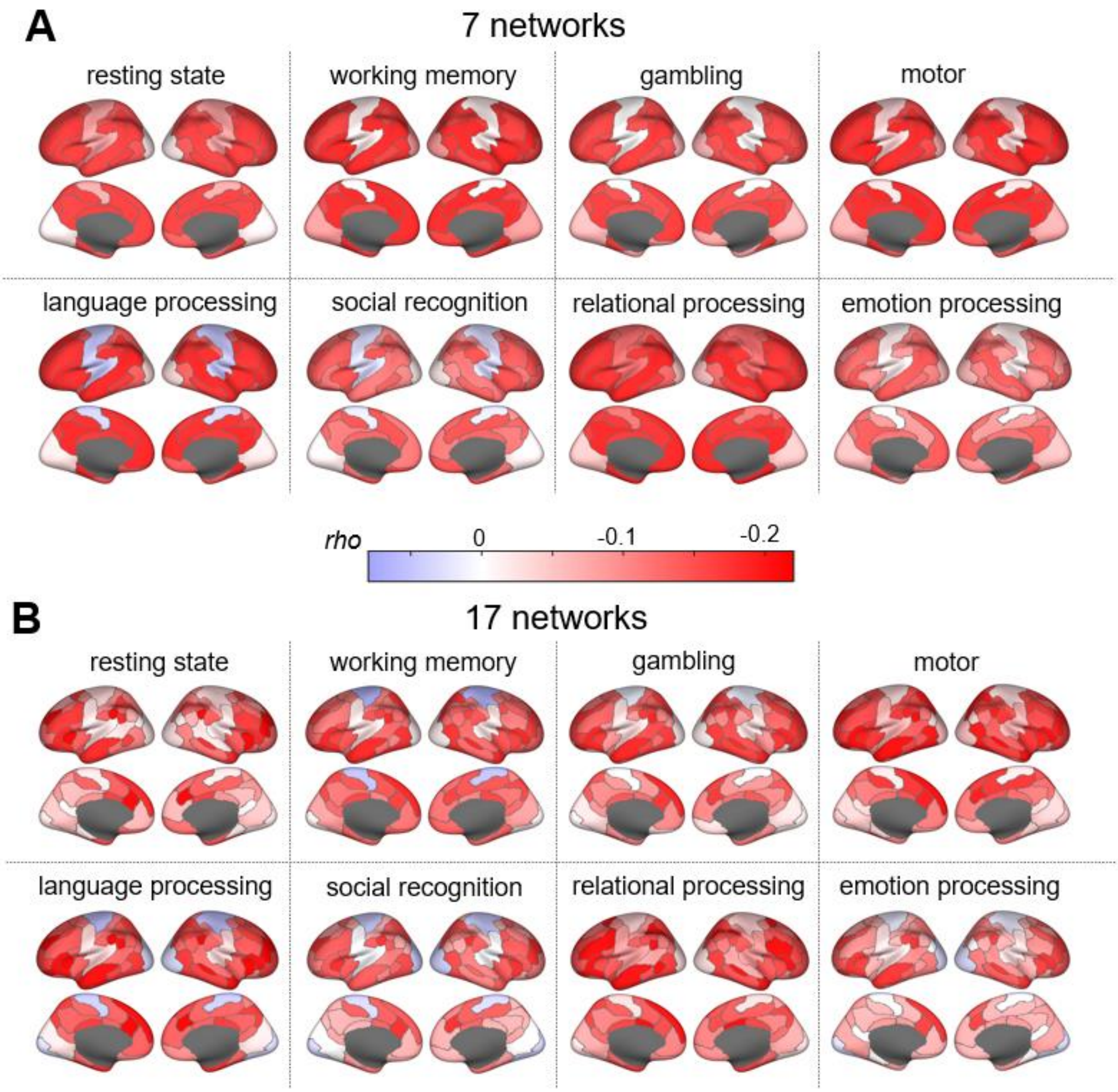


**Fig. S8.** Intelligence and state-specific brain network reconfiguration for seven (**A**) and 17 (**B**) functional brain networks. Note that the upper panel is similar to Fig. 2H of the main manuscript but shown here again for comparability. The strengths of Spearman correlations (*rho*) between intelligence (*g*-factor derived from 12 cognitive tasks) and brain network reconfiguration (cosine distance between functional connectivity matrices of different states) are illustrated in different colors (see color bar). Functional brain networks were derived from the Yeo atlas (Yeo et al. 2011). For calculating state-specific correlations, cosine distances were averaged over all state combinations a respective state was involved in (for task states, only combinations with different tasks were included). Similarly, the network-specific scores result from averaging over all network combinations (within and between network connectivity) in which the respective network was involved. All correlations were controlled for influences of age, sex, handedness, and in-scanner head motion (mean framewise displacement).

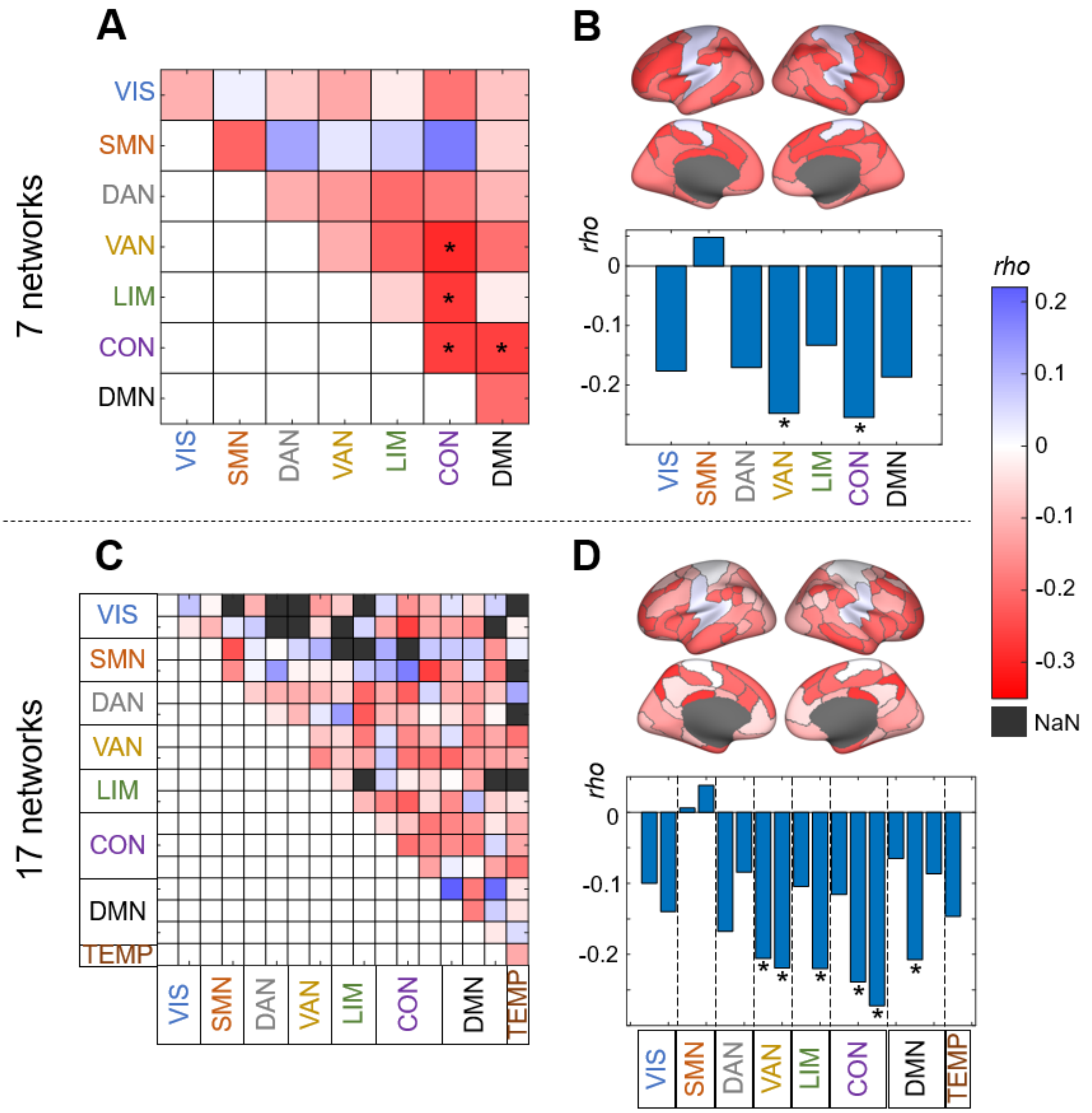


**Fig. S9.** Brain network-specific associations between intelligence and brain network reconfiguration for replication sample 1. Spearman correlation (*rho*) between intelligence (sum score of the Raven's Advanced Progressive Matrices Test, Raven et al. 1998) and brain network-specific reconfiguration values (averaged cosine distance between functional connectivity matrices of eight different cognitive states) for seven and 17 separate functional brain networks (Yeo et al. 2011). Network-specific correlations were projected onto the surface of the brain. The strengths of correlations are depicted in different colors (see color bar). All significant correlations (FDR-corrected *p*-values, $\alpha = .05$) are marked with asterisks. Note that NaN (not a number) values exist if in a specific network combination no single brain connection passes the filtering procedure (see Fig. S1 and Methods). (**A**) Associations between intelligence and brain network combination-specific reconfiguration scores for seven functional brain networks. (**B**) Associations between intelligence and reconfiguration scores for seven functional brain networks (averaged across all within- and between network combinations a respective network is involved in). (**C**) Associations between intelligence and brain network combination-specific reconfiguration scores for 17 functional brain networks. (**D**) Associations between intelligence and reconfiguration scores for 17 functional brain networks (averaged across all within- and between network combinations a respective network is involved in). All correlations were controlled for influences of age, sex, handedness, and in-scanner head motion (mean framewise displacement). FDR, false discovery rate; VIS, visual network; SMN, somatomotor network; DAN, dorsal attention network; VAN, salience/ventral attention network; LIM, limbic network; CON, control network; DMN, default mode network; TEMP, temporal parietal network.

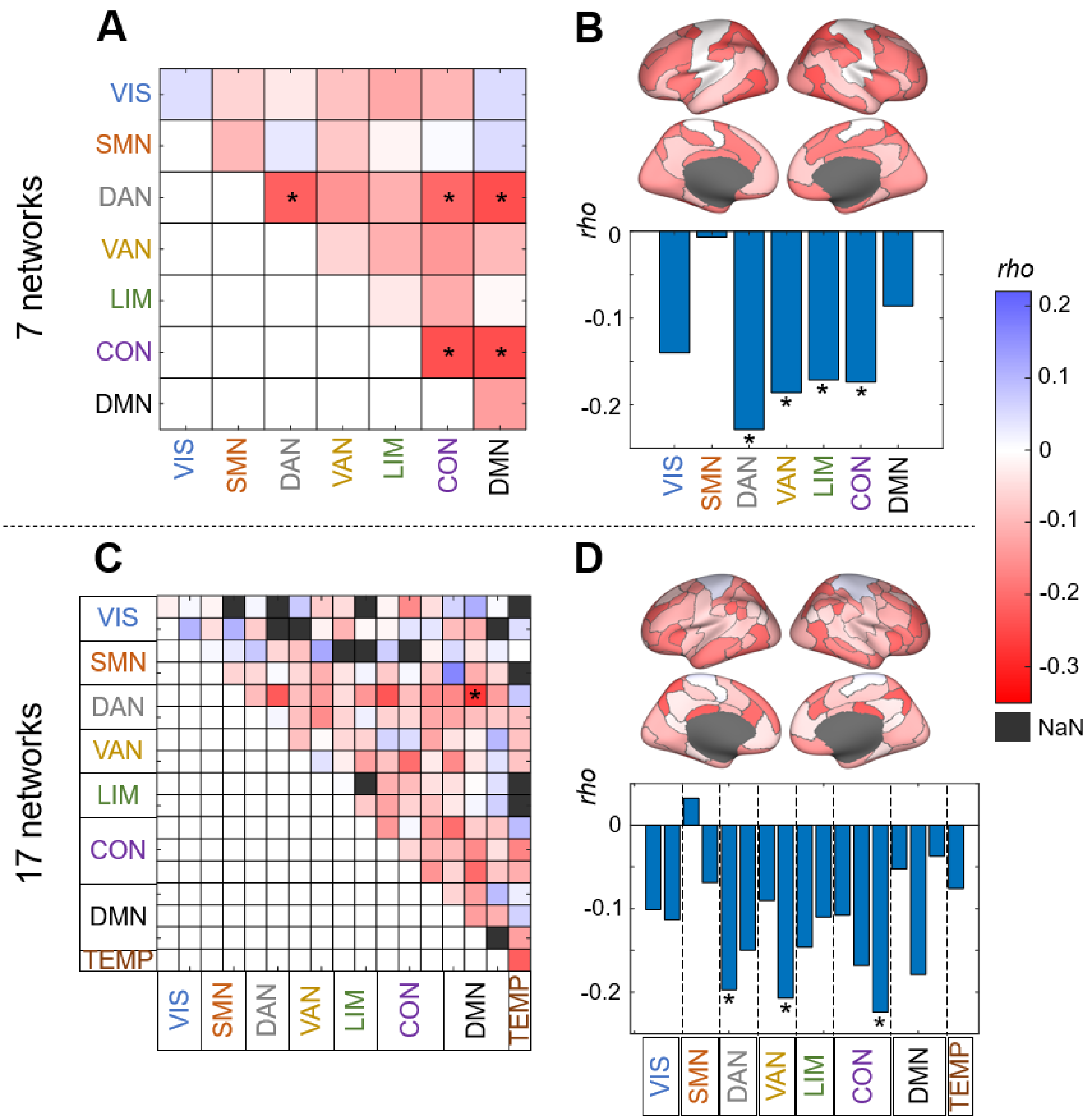


**Fig. S10.** Brain network-specific associations between intelligence and brain network reconfiguration for replication sample 2. Spearman correlation (*rho*) between intelligence (sum score of the Raven's Advanced Progressive Matrices Test, Raven et al. 1998) and brain network-specific reconfiguration values (averaged cosine distance between functional connectivity matrices of eight different cognitive states) for seven and 17 separate functional brain networks (Yeo et al. 2011). Network-specific correlations were projected onto the surface of the brain. The strengths of correlations are depicted in different colors (see color bar). All significant correlations (FDR-corrected *p*-values, $\alpha = .05$) are marked with asterisks. Note that NaN (not a number) values exist if in a specific network combination no single brain connection passes the filtering procedure (see Fig. S1 and Methods). (**A**) Associations between intelligence and brain network combination-specific reconfiguration scores for seven functional brain networks. (**B**) Associations between intelligence and reconfiguration scores for seven functional brain networks (averaged across all within- and between network combinations a respective network is involved in). (**C**) Associations between intelligence and brain network combination-specific reconfiguration scores for 17 functional brain networks. (**D**) Associations between intelligence and reconfiguration scores for 17 functional brain networks (averaged across all within- and between network combinations a respective network is involved in). All correlations were controlled for influences of age, sex, handedness, and in-scanner head motion (mean framewise displacement). FDR, false discovery rate; VIS, visual network; SMN, somatomotor network; DAN, dorsal attention network; VAN, salience/ventral attention network; LIM, limbic network; CON, control network; DMN, default mode network; TEMP, temporal parietal network.

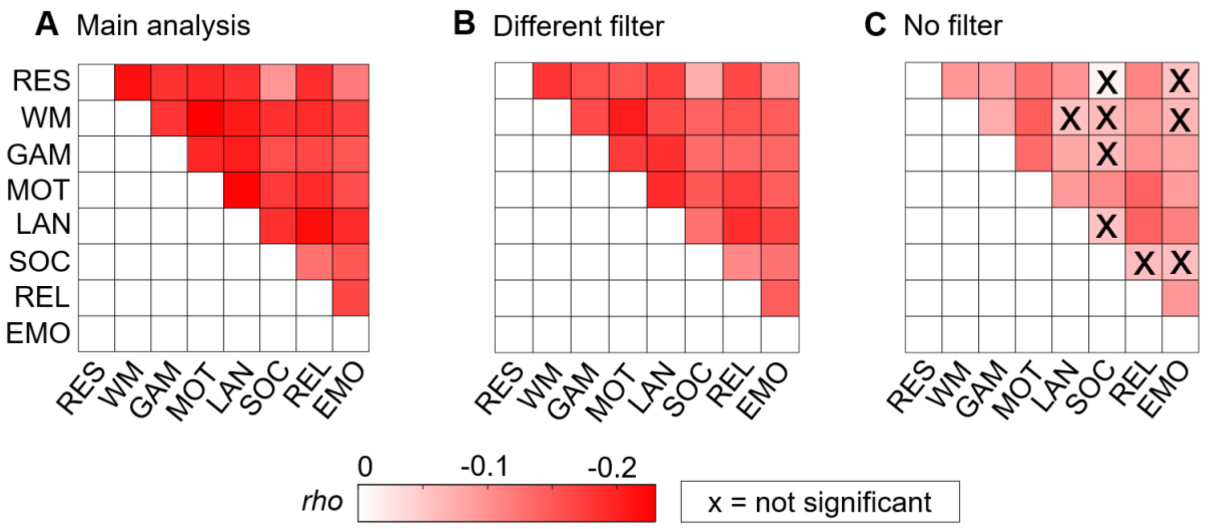


**Fig. S11.** Results of robustness control analyses for different filtering procedures of the functional connectivity matrices (FC). Spearman correlations (*rho*) between intelligence and brain network reconfiguration for all possible state combinations (FDR-corrected *p*-values, $\alpha$ = .05). The strengths of correlations are depicted in different colors (see color bar). All correlations were controlled for influences of age, sex, handedness, and in-scanner head motion (mean framewise displacement) and reached significance except the marked correlations. (**A**) Main analysis: FCs were filtered according to their correlation with intelligence ($p < .1$) and inconsistent connections, i.e., functional connections correlating in both positive and negative direction with intelligence across all states, were removed. (**B**) Different filter: FCs were filtered according to their correlation with intelligence ($p < .1$). (**C**) No filter. FDR, false discovery rate; RES, resting state; WM, working memory task; GAM, gambling task; MOT, motor task; LAN, language processing task; SOC, social cognition task; REL, relational processing task; EMO, emotion processing task.

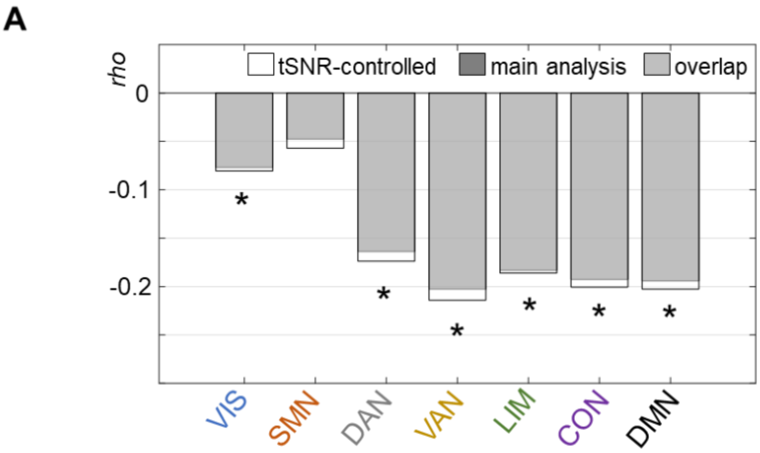


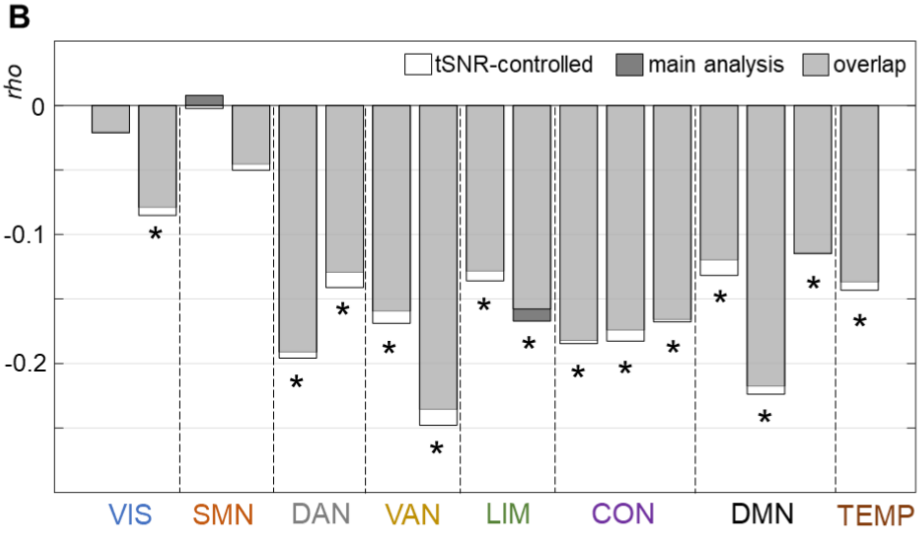


**Fig. S12.** Association between intelligence and brain network reconfiguration with additional control of brain network- and subject-specific variations in signal to noise ratio (tSNR). Associations are illustrated for seven (A), and 17 (B) brain networks (Yeo et al. 2011). Reconfiguration scores were averaged across all within- and between network combinations a respective network is involved in. The bar plots represent the comparison between results from the main analysis (Spearman correlations (*rho*) between intelligence and brain network reconfiguration controlled for age, sex, handedness, and in-scanner head motion, similar to Fig. 2E,G) and results from a post-hoc control analysis (Spearman correlations (*rho*) between intelligence and brain network reconfiguration controlled for age, sex, handedness, in-scanner head motion and tSNR; tSNR-controlled). Results from the main analysis are displayed in dark gray, results additionally controlled for tSNR in white, and the overlap between both in light gray. All significant correlations (FDR-corrected *p*-values, $\alpha$ = .05) are marked with asterisks. VIS, visual network; SMN, somatomotor network; DAN, dorsal attention network; VAN, salience/ventral attention network; LIM, limbic network; CON, control network; DMN, default mode network; TEMP, temporal parietal network.

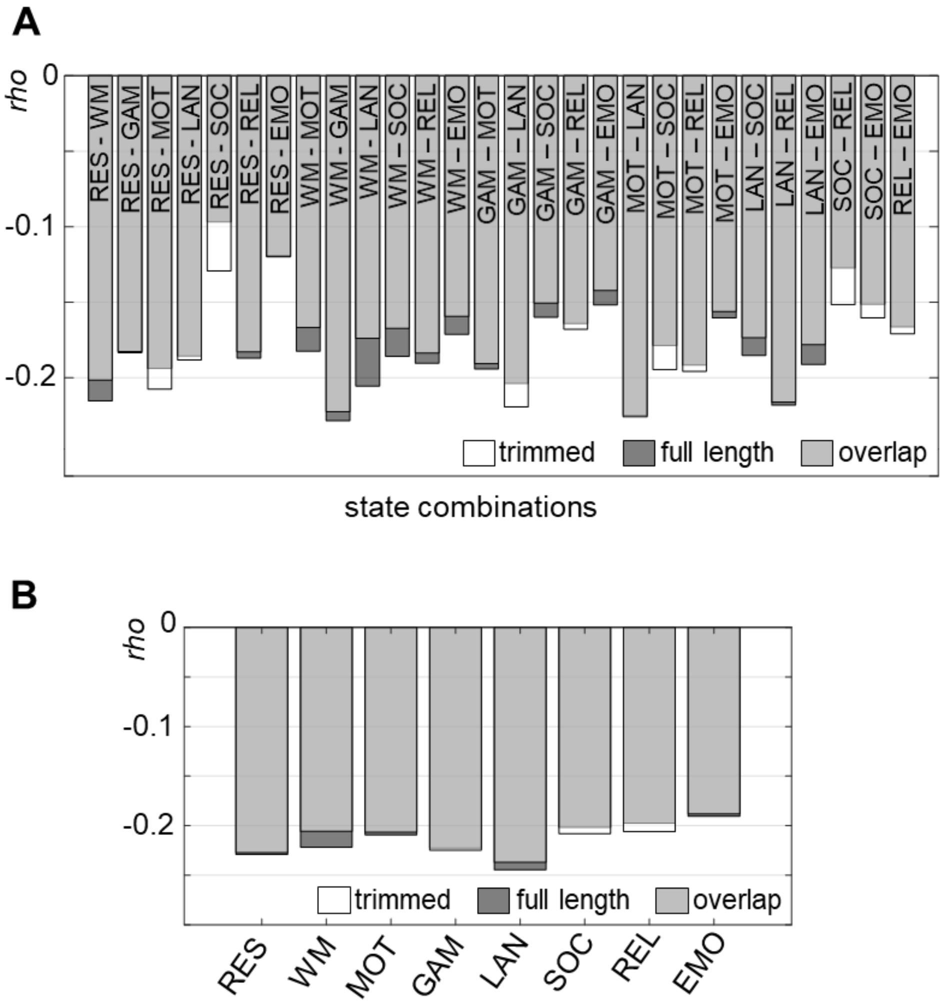


**Fig. S13.** Association between intelligence and brain network reconfiguration calculated on equally reduced scan length. Associations between intelligence and brain network reconfiguration calculated on the basis of the full lengths of each scan (depicted in dark gray) in contrast to the associations resulting from reduced scan lengths (176 consecutive timepoints each), depicted in white. The overlap between both is displayed in light gray. Each bar shows the Spearman correlations (*rho*) for both full and reduced scan length for one scan combination or specific state respectively. (**A**) Spearman correlations between intelligence and brain network reconfiguration for all possible state combinations. (**B**) Spearman correlations between intelligence and a total measure of state-specific reconfiguration. All correlations are significant (FDR-corrected *p*-values, $\alpha$ = .05). RES, resting state; WM, working memory task; GAM, gambling task; MOT, motor task; LAN, language processing task; SOC, social cognition task; REL, relational processing task; EMO, emotion processing task.
